# Structures of the human cholecystokinin receptors in complex with agonists and antagonists

**DOI:** 10.1101/2021.05.19.444887

**Authors:** Xuefeng Zhang, Chenglin He, Mu Wang, Qingtong Zhou, Dehua Yang, Ya Zhu, Wenbo Feng, Hui Zhang, Antao Dai, Xiaojing Chu, Jia Wang, Zhenlin Yang, Yi Jiang, Ulrich Sensfuss, Qiuxiang Tan, Shuo Han, Steffen Reedtz-Runge, Eric H. Xu, Suwen Zhao, Ming-Wei Wang, Beili Wu, Qiang Zhao

## Abstract

Cholecystokinin receptors, CCK_A_R and CCK_B_R, are important neuro-intestinal peptide hormone receptors and play a vital role in food intake and appetite regulation. Here we report three crystal structures of the human CCK_A_R in complex with different ligands, including one peptide agonist and two small-molecule antagonists, as well as two cryo-electron microscopy structures of CCK_B_R–gastrin in complex with G_i2_ and G_q_, respectively. These structures reveal the recognition pattern of different ligand types and the molecular basis of peptide selectivity in the cholecystokinin receptor family. By comparing receptor structures in different conformational states, a stepwise activation process of cholecystokinin receptors is proposed. Combined with pharmacological data, our results provide atomic details for differential ligand recognition and receptor activation mechanisms. These insights will facilitate the discovery of potential therapeutics targeting cholecystokinin receptors.

---

Biologically active peptides often present in families whose members display sequence and structural similarity. Cholecystokinin (CCK) and gastrin, however, are the only two members of the dityrosyl-sulfated peptide family that exists in mammals and share the same carboxyl-terminal octapeptide-amide^1, 2^. They are the most abundant peptides in the gastrointestinal tract and the central nervous system acting as physiologically important hormones and neurotransmitters. Among CCK and gastrin peptides of different lengths, CCK-8 and gastrin-17 are the major forms with full biological activity^3–5^. Cholecystokinin A and cholecystokinin B receptors (CCK_A_R and CCK_B_R) are the two homologous G protein-coupled receptors (GPCRs) for CCK and gastrin, respectively^6, 7^. CCK_A_R preferentially binds to sulfated CCK, while CCK_B_R recognizes both CCK and gastrin with similar affinities and discriminate poorly between sulfated and non-sulfated forms of CCK and gastrin^7–9^. Previous studies demonstrated that gastrin could activate pertussis toxin-sensitive or phospholipase C-dependent downstream effectors, indicating both G_i/o_ and G_q/11_ signaling are involved in CCK_B_R functionality^10, 11^. When activated, these two receptors engage in fundamental physiological actions such as satiety regulation, pancreatic enzyme secretion and gall bladder contraction^6, 12^. They are also implicated in behavioral processes including anxiety, memory and drug addiction^13, 14^.

In line with the important roles of CCKRs, their ligands have shown therapeutic potential for anxiety, obesity and type 2 diabetes^15, 16^. Highly selective non-peptidic antagonists of CCK_A_R such as devazepide [3S(-)-N(2,3-Dihydro-1-methyl-2-oxo-5-phenyl-1H-1,4-benzodiazepine-3-yl)-1H-indole-2-carboxamide] and lintitript [1-([2-(4-(2-Chlorophenyl) thiazole-2-yl)aminocarbonyl]indolyl) acetic acid] (Extended Data Fig. 1) were developed to treat gastrointestinal disorders, neuropathic pain and pancreatic cancer^17, 18^, while highly selective CCK_A_R agonists such as NN9056, a modified CCK-8 with high selectivity and long-acting properties, have been proposed as potential treatment for obesity^19, 20^. Meanwhile, antagonists such as Z-360 or JB95008 and agonists such as ceruletide targeting CCK_B_R have been implicated in treating diabetes, anxiety and thyroid cancer, etc^21, 22^. In addition, antagonists of CCKRs, such as devazepide for CCK_A_R or L365,260 for CCK_B_R, has exhibit the significant attenuation in drug or stress induced relapse to cocaine seeking in animal model, showing potential in treating drug abuse^23^. However, many clinical trials targeting CCK_A_R or CCK_B_R were terminated at different phases due to low efficacy or poor bioavailability in patients, suggesting that a better understanding of the CCKR family is required. To provide molecular details of ligand recognition and receptor activation of CCKRs, we solved the crystal structures of the human CCK_A_R in complex with two small-molecule antagonists (lintitript and devazepide) and one full agonist NN9056, as well as two cryo-electron microscopy (cryo-EM) structures of CCK_B_R in complex with gastrin coupled to G_i_ and G_q_, respectively.

## Overall structures of CCKR complexes

To improve protein stability and facilitate crystallization, residues K241–S301 in the third intracellular loop (ICL3) of CCK_A_R were replaced with a T4 lysozyme fusion protein and 22 residues at the receptor C terminus were truncated. Additionally, a mutation F130^3.41^W (superscript indicates nomenclature according to Ballesteros-Weinstein numbering system^24^) was introduced to further improve protein homogeneity. Using the optimized CCK_A_R, the CCK_A_R–NN9056 complex structure was determined at 3.0 Å resolution (Extended Data Table 1). To solve the structure of CCK_A_R bound to the antagonists, another mutation D87^2.50^N was introduced to stabilize the receptor in an inactive state^25^, and the complex structures of CCK_A_R–devazepide and CCK_A_R–lintitript were determined at 2.5 Å and 2.8 Å resolution, respectively (Extended Data Fig. 2, Extended Data Table 1).

The CCK_A_R structures adopt a canonical seven-transmembrane helical bundle structure (helices I-VII) of GPCRs (Fig. 1a–d), with the second extracellular loop (ECL2) of CCK_A_R forming a β-hairpin structure that is similar to the previously determined structures of peptide receptors^26–28^. The third extracellular loop (ECL3) adopts a two-turn α-helical conformation which has never been observed before in other class A GPCR structures. The complexes of CCK_A_R–devazepide and CCK_A_R–lintitript are structurally similar with an overall Cα root-mean-square deviation (RMSD) of 0.33 Å (Fig. 1g, h). Compared to the CCK_A_R–lintitript structure, the CCK_A_R–NN9056 structure showed a larger variance (overall Cα RMSD, 0.48 Å) with main differences appearing in the ligand binding region of helix VI. Upon binding to the agonist NN9056, ~1 Å inward shift of helix VI in the ligand binding region was observed and such a conformational change might be required for receptor activation. Except for helix VI, the differences in the transmembrane domain between agonist- and antagonist-bound CCK_A_R are relatively small.

**Fig. 1.**
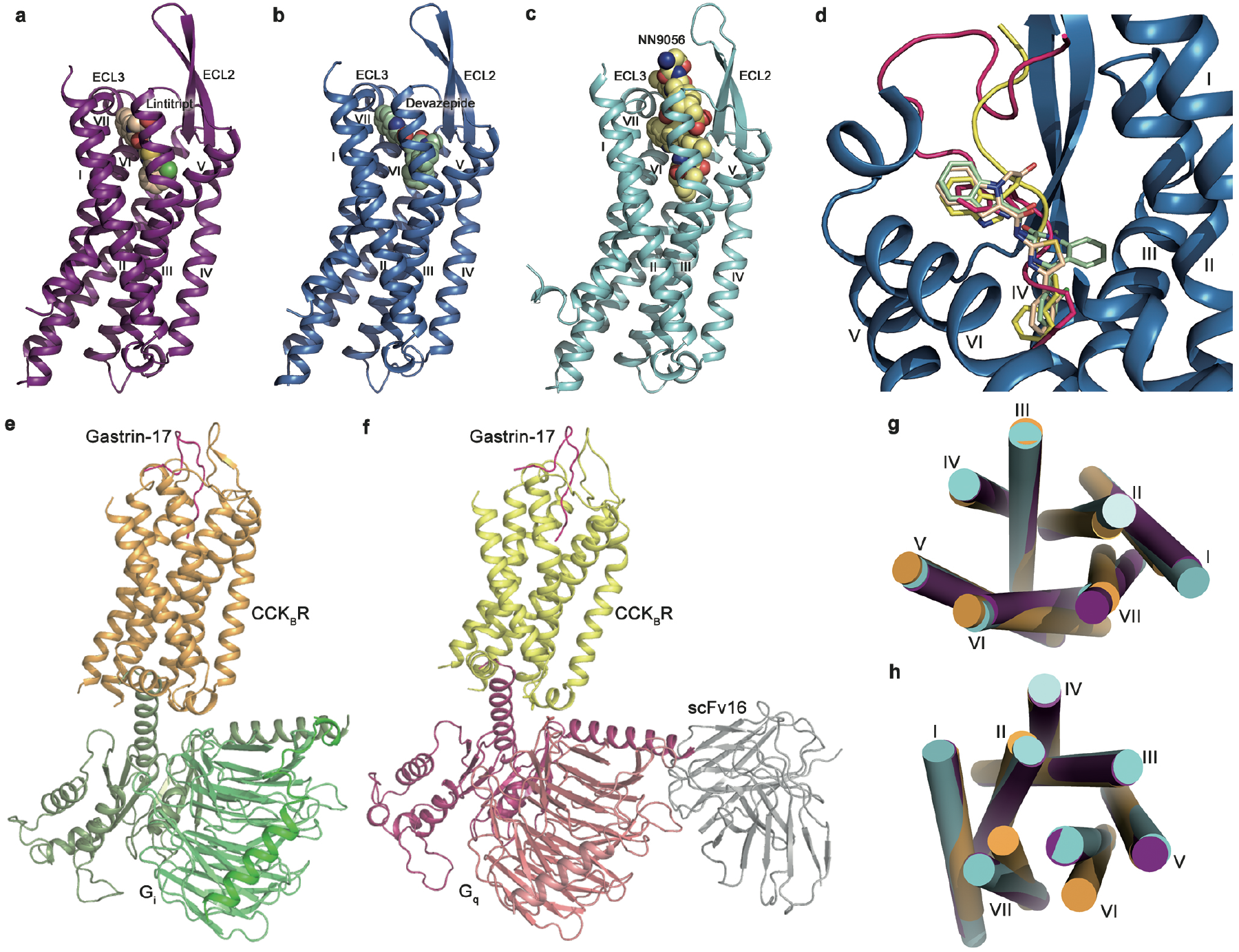
Overall structures of the CCK_A_R and CCK_B_R. **a–c**, Crystal structures of the CCK_A_R–lintitript (**a**), CCK_A_R–devazepide (**b**) and CCK_A_R–NN9056 (**c**) complexes. The receptors are shown in deep-purple, marine and cyan cartoon representation, respectively. Lintitript is shown as spheres with carbons in wheat. Devazepide is shown as spheres with carbons in pale-green. NN9056 is shown as spheres with carbons in pale-yellow. **d**, the comparison of antagonists and agonists binding position superimposed in CCK_A_R. The antagonists are shown in sticks and peptide agonists are shown in cartoon with key residues in stick. All the ligands are colored as in corresponding panel in this figure. **e, f**, Cryo-EM structures of CCK_B_R in complex with gastrin-17 and G_i_ or G_q_. The receptors are shown in bright-orange and yellow, respectively. Gastrin-17 in shown as hot-pink sticks. The G_i2_ trimers are shown in smudge, lime and green cartoons, and the G_q_ trimers are shown in warm-pink, deep-salmon and salmon cartoons, respectively. **g**, **h**, Comparison of helical bundles of CCK_A_R in inactive (CCK_A_R–lintitript), partially active (CCK_A_R–NN9056) and CCK_B_R in fully active state (gastrin-17–CCK_B_R–Gi) on extracellular (**f**) and intracellular (**g**) sides. The receptors are colored as described above.

It was reported that CCK_B_R could activate both G_i_ and G_q_ signaling pathways, and indeed, CCK_B_R was co-purified with these two G proteins while failed to form protein complex with Gs. The CCK_B_R–G_q_ and CCK_B_R–G_i2_ complexes were assembled in the presence of the CCK_B_R selective endogenous peptide gastrin-17 and their structures were determined by single-particle cryo-EM analysis with the overall resolution of 3.1 Å and 3.3 Å (Fig. 1e, f, Extended Data Fig. 3, Extended Data Table 2), respectively. Compared to the inactive CCK_A_R structure, the G_q_/G_i2_-coupled CCK_B_R structures exhibit key structural features of active class A GPCRs, including a large outward movement of the intracellular tip of helix VI and a small inward shift at the intracellular end of helix VII. However, the conformational changes in CCK_B_R are relatively small with an approximately 6 Å movement for helix VI compared to other active structures of class A GPCR–G-protein complexes^29–31^.

These two G protein-coupled CCK_B_R complexes are structurally similar with an overall Cα RMSD of 0.9 Å. However, despite the overall similarity, several distinct features have been observed in the recognition patterns for G_i2_ and G_q_. Both G_i2_ and G_q_ insert in the same binding pocket formed by helices II, III, VI and VII as well as the intracellular loops, however, a ~8°rotation between the G_i2_ and G_q_ heterotrimeric proteins was noted, as described in the companion CCK_A_R paper (Extended Data Fig. 4). Comparison of the structures of CCK_B_R–G_i2_ and CCK_B_R–G_q_ complexes revealed differences in the intracellular loops and transmembrane domain of CCK_B_R as well as the αN helix and C-terminal α5 helix of G protein. The distance between the receptor ICL2 and the αN helix of the CCK_B_R–G_q_ complex is closer than that in the CCK_B_R–G_i2_ complex, resulting in more extensive interactions between ICL2 of CCK_B_R and the αN helix of G_q_. R163^ICL2^ and V164^ICL2^ in CCK_B_R form several hydrophobic interactions with the αN helix of G_q_ while only V164^ICL2^ forms hydrophobic interaction with that of G_i2_. In addition, Q166^ICL2^ and T167^ICL2^ form several hydrogen bonds with G_q_ while only one hydrogen bond is formed between ICL2 of CCK_B_R and G_i2_.

Since the two G protein-coupled CCK_B_R complexes are highly resemble to each other, the CCK_B_R–G_i2_ complex was used to compare with the corresponding complex structure of CCK_A_R and these structures are very similar with an overall Cα RMSD of 0.9 Å. The CCK_A_R and CCK_B_R complexes showed a common activation pattern with W^6.48^ and the PIF motif exhibiting similar confirmations upon activation. The G_i2_ binds to CCK_A_R and CCK_B_R in a similar binding pocket formed by helices II, III and V-VIII. The outward movement of the intracellular tip of helix VI and inward shift at the intracellular end of helix VII of the two CCKRs are within a similar range, suggesting that these receptors activate G proteins by the same mechanism.

## Binding modes of small molecules in CCK_A_R

Devazepide binds to CCK_A_R in a pocket bordered by helices II-VII, ECL2 and ECL3 (Fig. 2a). This antagonist is characterized by a 1,4-benzodiazepine group and an indole moiety^32^. Residue R336^6.58^ is engaged in a π–cation interaction with the indole group and forms a hydrogen bond with the carbonyl oxygen of the 1,4-benzodiazepine group (Fig. 2b). In addition, the 4-nitrogen within the benzodiazepine group and the amide nitrogen form two hydrogen bonds with the residue N333^6.55^. The critical role of N333^6.55^ and R336^6.58^ in devazepide binding was also reflected by a notable loss of antagonistic activity of devazepide in our NN9056-induced inositol phosphate (IP) accumulation assay (Extended Data Table 3) and a complete abolishment of CCK-8 binding ability for the mutants N333^6.55^A and R3 3 6^6.58^A (Extended Data Table 4). The hydrogen bond between devazepide and N333^6.55^ is important for its affinity, as previous studies showed that chiral change of the 3-carbon within the benzodiazepine group which eliminates the hydrogen bond led to an over 100-fold decrease of the binding affinity^33, 34^. In addition to the above interactions, the indole group of devazepide is further stabilized by hydrophobic interactions with residues A343^ECL3^, E344^ECL3^, L347^ECL3^, and I352^7.35^ in ECL3 and helix VII. The phenyl ring of the benzodiazepine group forms multiple hydrophobic interactions with N98^2.61^, T117^3.28^, and T118^3.29^ in helices II and III, while the tolyl group penetrates deeply into the binding pocket, making hydrophobic contacts with M121^3.32^, Y176^4.60^, and F330^6.52^. The roles of these residues in devazepide recognition were investigated by testing the effects of their alanine mutations on the antagonistic activity of devazepide. The results show that all these mutations impaired the inhibitory activity of devazepide on NN9056-induced IP production with the exceptions of N98^2.61^A and Y176^4.60^A (Extended Data Table 3). Among these mutations, T117^3.28^A and F330^6.52^A exhibited the largest effect, suggesting that they play a vital role in recognizing devazepide.

**Fig. 2.**
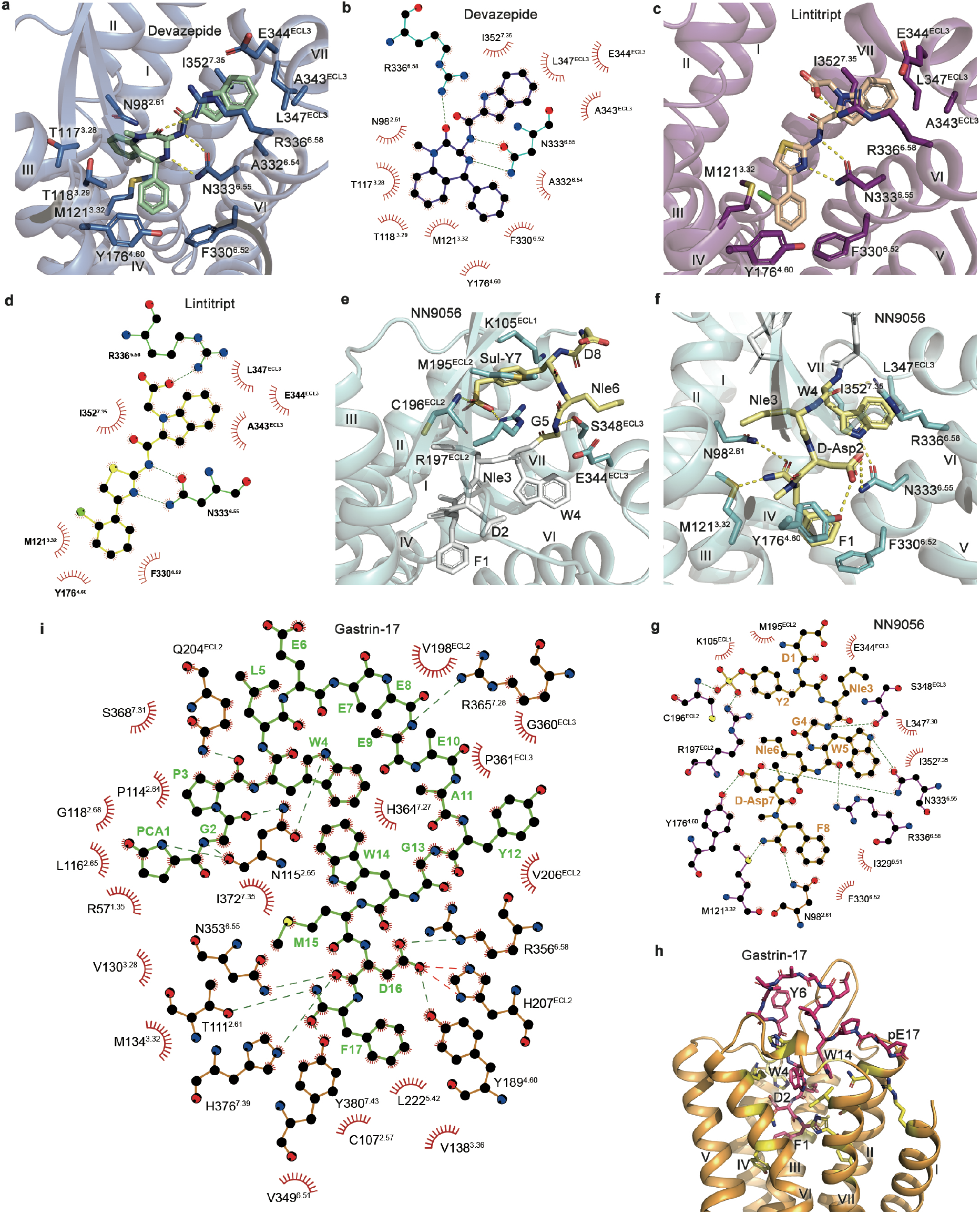
Small-molecule ligand binding pocket of CCKRs. **a**, Interactions between devazepide and CCK_A_R. The CCK_A_R residues involved in interactions are shown as cyan sticks. Devazepide is displayed as blue sticks. Polar interactions are shown as yellow dashed lines. **b**, Schematic representation of the interactions between CCK_A_R and devazepide analyzed using the LigPlot^+^ program^41^. Interactions between lintitript and CCK_A_R (**c**) and schematic representation (**d**). The CCK_A_R residues involved in interactions are shown as green sticks. Lintitript is displayed as yellow sticks. **e–g**, Interactions between top region (**e**) and bottom region (**f**) of NN9056 and CCK_A_R and schematic representation (**g**). The CCK_A_R residues involved in interactions are shown as brown sticks. NN9056 is displayed as magenta sticks. Polar interactions are shown as yellow dashed lines. Interactions between gastrin-17 and CCK_B_R (**h**) and schematic representation (**i**). The CCK_B_R residues responsible for important interactions are shown as pale-cyan sticks. Gastrin-17 is displayed as hot-pink sticks.

The antagonist lintitript occupies a binding pocket similar to devazepide (Fig. 2c, d). Instead of the bulky benzodiazepine group, lintitript has a thiazolyl group in the middle portion, limiting its interaction with helices II and III of the receptor and resulting in a lower binding affinity to CCK_A_R compared to that of devazepide^34^. Similar hydrogen bonds between lintitript and N333^6.55^ as well as π–cation interaction with R336^6.58^ were formed as observed in the CCK_A_R–devazepide structure, despite their different chemical scaffolds. These polar interactions are also critical for lintitript recognition, as alanine replacements of N333^6.55^ and R336^6.58^ reduced its antagonistic activity by 6-fold and 8-fold, respectively, and completely abolished the binding of CCK-8. Similar to the indole and tolyl groups of devazepide, the corresponding groups in lintitript also form multiple hydrophobic interactions with the receptor. The alanine mutations of the residues that are involved in these hydrophobic interactions diminished the antagonistic effect of lintitript, except for Y176^4.60^A (Extended Data Table 3).

## Binding modes of peptide ligands in CCK_A_R and CCK_B_R

The full CCK_A_R agonist NN9056 is a highly selective peptide analogue synthesized based on CCK-8^19^. It was developed mainly by introducing D-N-methyl-Asp and N-methyl-Phe instead of Asp and Phe at the penultimate position and replacing Met with Nle in CCK-8 as well as adding a C18-acylated fatty chain to the N terminus. Unambiguous electron densities were observed for the C-terminal octapeptide in NN9056, while the long N-terminal modification of the agonist was not modeled due to the poor density map. NN9056 adopts a binding pose perpendicular to the membrane plane, with its N terminus pointing to the extracellular surface and the C terminus penetrating into the helical bundle (Fig. 2e, f). The N terminus of NN9056 is anchored to ECL2 through a salt bridge between the sulfonate group of Y7 (as counted from the C terminus of CCK; the residues in gastrin is numbered in the same way) and residue R197 in CCK_A_R, which is crucial for CCK_A_R selectivity as the nonsulfated CCK binds selectively to CCK_B_R^35^. Besides R197, the NN9056 residue Y7 also makes hydrophobic contacts with residues K105 and M195 in ECL1 and ECL2. This aligns well with previous studies suggesting that M195^ECL2^ is a binding partner for the sulfated tyrosine of CCK^36^. The extracellular binding moiety of NN9056 is further stabilized by a hydrogen bond between the main-chain nitrogen of G5 and the side chain of S348 in ECL3. The side chain of R197^ECL2^, which forms an important salt bridge with the sulfated Y7, is only 3.4 Å away from the Cα of G5. Thus, any other residues replacing this glycine will import a bulkier side chain, which would cause steric clash with R197 and reduce the binding of CCK-8. Indeed, it was reported that all mutations on G5, with the exception of N-methylglycin, led to significantly reduced binding affinities^19^.

The C-terminal region of NN9056 occupies a similar binding site to that of devazepide and lintitript, with the side chains of residues W4 and F1 in the agonist overlapping with the indole and tolyl groups of the antagonists. The side chain of W4 extends toward helices VI and VII, and ECL3, forming a hydrogen bond with N333^6.55^ and hydrophobic interactions with L347^7.30^ and I352^7.35^, while the NN9056 residue D2 builds two hydrogen bonds with Y176^4.60^ and N333^6.55^ (Fig. 2f, g). Additionally, the side chain of F1 interacts with a hydrophobic cluster formed by Y176^4.60^, I329^6.51^, and F330^6.52^ in CCK_A_R. The C-terminal amide of NN9056 makes two hydrogen bonds with N98^2.61^ and M121^3.32^ to further stabilize the binding between the C terminus of NN9056 and the receptor. These two polar interactions were supported by mutagenesis studies showing that alanine replacements of N98^2.61^ and M121^3.32^ were associated with an over 2-fold reduction of NN9056 potency in inducing IP production. Different from the mutagenesis results of CCK_A_R with NN9056, the mutations N98^2.61^A, Y176^ECL2^A, N333^6.55^A and R336^6.58^A displayed a much more predominant influence showing a 10–200-fold reduction of CCK-8 potency in inducing receptor activation, probably due to the fact that modifications in NN9056 stabilize the peptide (Extended Data Table 4). Additionally, two Nle residues in NN9056 do not form any strong interaction with the receptor, which explains why replacing them with methionine or leucine did not affect the affinity^37, 38^.

Unlike the linear peptides CCK and NN9056, the 17-amino-acid peptide gastrin forms a β-hairpin structure (Fig. 2h, i). The five residues in the C terminus insert deeply into the ligand-binding pocket formed by helices II, III, V, VI, and VII, as well as ECL2 and ECL3, a binding site similar to that in the crystal structure of CCK_A_R–NN9056. The three amino acids at the C terminus occupy a similar site to that for the side chain of the sulfated tyrosine of NN9056 in CCK_A_R (Fig. 3c). This binding mode is supported by our mutagenesis assays, in which alanine mutations of Y189^4.60^, R356^6.55^, L367^ECL3^ and Y380^7.43^ completely abolished the binding of CCK-8 or gastrin-17 (Extended Data Table 6), suggesting key roles of these residues in peptide recognition. In addition to the interactions engaged by the peptide C terminus, residues at the N terminus of gastrin-17 locate in a shallow binding cavity shaped by ECL1, ECL2 and ECL3 further improve the binding affinity and ligand selectivity (Fig. 3a, b).

## Peptide selectivity between CCK_A_R and CCK_B_R

Both gastrin and CCK are vertebrate brain-gut peptides that share a conserved common C-terminal pentapeptide amide sequence^4, 5^ crucial for biological functions, while the N-terminal extensions, especially the tyrosine residue at position 7 of CCK and position 6 of gastrin (as counted from the peptide C terminus phenylalanine), serve to increase potency and specificity. Unlike sulfation of the residue Y6 in gastrin, sulfation of CCK-8 in Y7 is critical to its affinity at CCK_A_R^39^. The residues in the binding pockets of CCK_A_R and CCK_B_R are highly conserved, however, minor amino acid differences between the two receptors reshape their binding pockets and allow different peptide preference.

As previously described, the C termini of NN9056 and gastrin bind to their respective receptors in a similar manner and the detailed interactions between the three C-terminal residues of the peptide and the receptor are almost identical (Fig. 3c). The main difference in this region is that the histidine residue at position 7.39 of CCK_B_R forms a hydrogen bond with the main-chain carbonyl of D2 in gastrin, which does not exist in the CCK_A_R–NN9056 complex as the counterpart in CCK_A_R is a leucine. However, starting from W4, these two peptides show different orientations and thus lead to different binding modes. This is due to sequence variety in ECL2 between the CCKR family: (i) the residue L200^ECL2^ at the end of ECL2 in CCK_A_R is substituted by W209^ECL2^ in CCK_B_R, where the bulky side chain pushes the side chain of H207^ECL2^ toward gastrin to enable a hydrogen bond with the main chain of W4, dragging this peptide further toward ECL2 in comparison with NN9056 (Fig. 3c). This is supported by our mutagenesis data showing that replacing H207^ECL2^ with an alanine completely abolished the gastrin binding (Extended Data Table 5); (ii) the key residue R197^ECL2^ in CCK_A_R, which forms a key salt bridge with the sulfated tyrosine in NN9056, is not conserved and appears as a valine in CCK_B_R. The long side chain of R197^ECL2^ is either locked by this salt bridge or by the negatively charged residue E344^ECL3^, and would cause a severe spatial hindrance with gastrin and does not allow this peptide to bind to CCK_A_R in the same manner (Fig. 3e, f). Therefore, our structures explain why gastrin could only bind to CCK_A_R with a very weak affinity in contrast to CCK. The importance of this polar interaction was confirmed by our mutagenesis studies, in which the CCK_A_R mutant R197^ECL2^A greatly diminished the binding of CCK-8 and reduced the agonist potency of NN9056 by about 3-fold in the IP accumulation assay (Extended Data Table 5).

The above differences in the binding modes of the peptide agonists result in a distinct binding environment for the sulfated tyrosine residue. For NN9056, the sulfonate group of Y7 (sul-Y7) anchors into a binding cavity formed by helix II, ECL1 and ECL2 of CCK_A_R (Fig. 3g). The salt bridge between the sul-Y7 and R197^ECL2^ breaks the original salt bridge between R197^ECL2^ and E344^ECL3^, opens up the binding pocket, and allows the entrance of CCK (Fig. 3d, g). This is consistent with the fact that the sulfation of CCK improves its affinity by 1,000 folds towards CCK_A_R^40^. In the case of CCK_B_R, however, the lack of the spatial hindrance of R197^ECL2^ side chain allows gastrin to bind in a more extended manner, and thus, the gastrin residue Y6 occupies a position similar to that of the NN0956 residue Y7 but rotated by about 90°, only making hydrogen bonds with ECL2 (Fig. 3b, h). One non-conserved residue, R208^ECL2^ in CCK_B_R (L199^ECL2^ in CCK_A_R), reaches out to form a hydrogen bond with the hydroxyl group of Y7 resulting in an increased peptide affinity. It could be expected that sulfation of this tyrosine would form electrostatic interactions with R208^ECL2^ and further improve the affinity of gastrin to some extent and indeed, previous studies have shown that the sulfated gastrin displayed a 10-fold higher binding affinity compared to the non-sulfated form^39^.

The N-terminal extension of gastrin, especially the tryptophan and pyroglutamic acid residues at positions 14 and 17, also interacts with the receptor through several hydrogen bonds with residues N115^2.65^ and R57^1.35^ in CCK_B_R to improve its affinity and selectivity. For instance, the side chain of W14 would form a severe clash with the side chain of R197^ECL2^ in CCK_A_R, thereby reducing the ability of gastrin to bind this receptor. In addition to the interactions with CCK_A_R, W14 also forms extensive interactions with W4 within gastrin, stabilizing the peptide in a conformation different from that of NN9056 (Fig. 3e, f).

## Stepwise activation process of CCK_A_R

CCK_A_R could be activated by CCK-8 both in sulfated and non-sulfated forms at different potencies. To study the binding mode of endogenous agonist CCK-8ns (non-sulfated CCK-8), docking and molecular dynamic simulation studies on the basis of the cryo-EM structure of CCK-8–CCK_A_R–G_q_ complex structure reported in our companion paper were performed (Fig. 4a). The predicted binding site of CCK-8ns in CCK_A_R is similar to the binding site of NN9056, which is a peptide mimic of the sulfated CCK-8, despite lacking of key interactions between the sulfonate group of NN9056 and CCK_A_R. Superimposing the CCK_A_R structures reveals that the NN9056-bound CCK_A_R is in a similar conformation to that of devazepide-bound receptors (Cα RMSD = 0.48), but differs from that of the G protein-coupled CCK_A_R reported in our companion paper (Cα RMSD = 1.86 Å) (Extended Data Fig. 5), suggesting that the agonist itself is not sufficient to fully stabilize the active conformation.

**Fig. 3.**
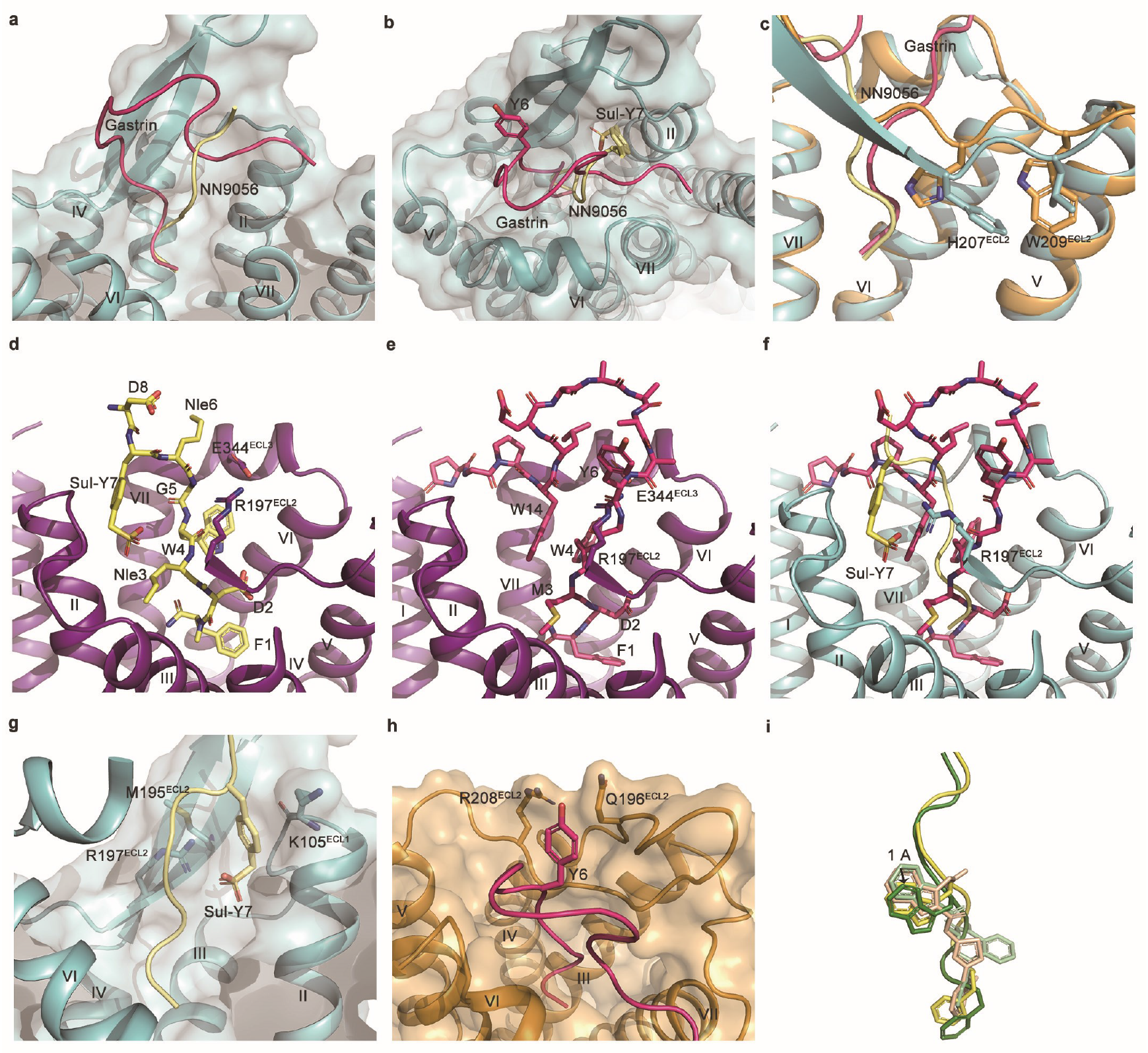
Comparison of peptide binding in CCKRs. **a**, Comparison of peptide conformation. The CCK_A_R and NN9056 are shown in cyan and pale-yellow cartoons, respectively. The gastrin-17 is shown in hot-pink cartoons and aligned to the same position in CCK_A_R.**b**, Comparison of important tyrosine residue in NN9056 and gastrin-17. The side chains of Y7 in CCK-8 and Y6 in gastrin-17 are shown in pale-yellow sticks and hot-pink sticks, respectively. **c**, Comparison of peptide C terminus in corresponding receptors. CCK_A_R and CCK_B_R are shown in cyan and bright-orange cartoons, respectively. NN9056 and gastrin-17 are shown in pale-yellow and hot-pink cartoon representation. The side chains of H207 and W209 of CCK_B_R and their corresponding residues in CCK_A_R are shown in sticks. **d**, Superposition of NN9056 in inactive CCK_A_R (represented by the CCK_A_R–lintitript structure). The receptor and NN9056 are shown in deep-purple cartoons and pale-yellow sticks, respectively. The salt bridge of R197 and E344 are shown in brown sticks. **e**, **f**, Superposition of gastrin-17 in inactive CCK_A_R (represented by the CCK_A_R–lintitript structure) and active CCK_A_R (represented by the CCK_A_R–NN9056 structure). The receptor and gastrin-17 are shown in deep-purple cartoons and hot-pink sticks, respectively. **g**, **h**, Peptide binding pocket of CCK_A_R–NN9056 and CCK_B_R–gastrin-17. The receptors are shown in cartoon and surface representation. NN9056 and gastrin-17 are shown in pale-yellow and hot-pink sticks, respectively. **i**, The binding position comparison of peptide agonists and small-molecule antagonists. The agonists NN9056 and CCK-8 are shown in pale-yellow and forest green cartoon and sticks, and small-molecule antagonists lintitript and devazepide are shown in wheat and pale-green sticks, respectively.

**Fig. 4.**
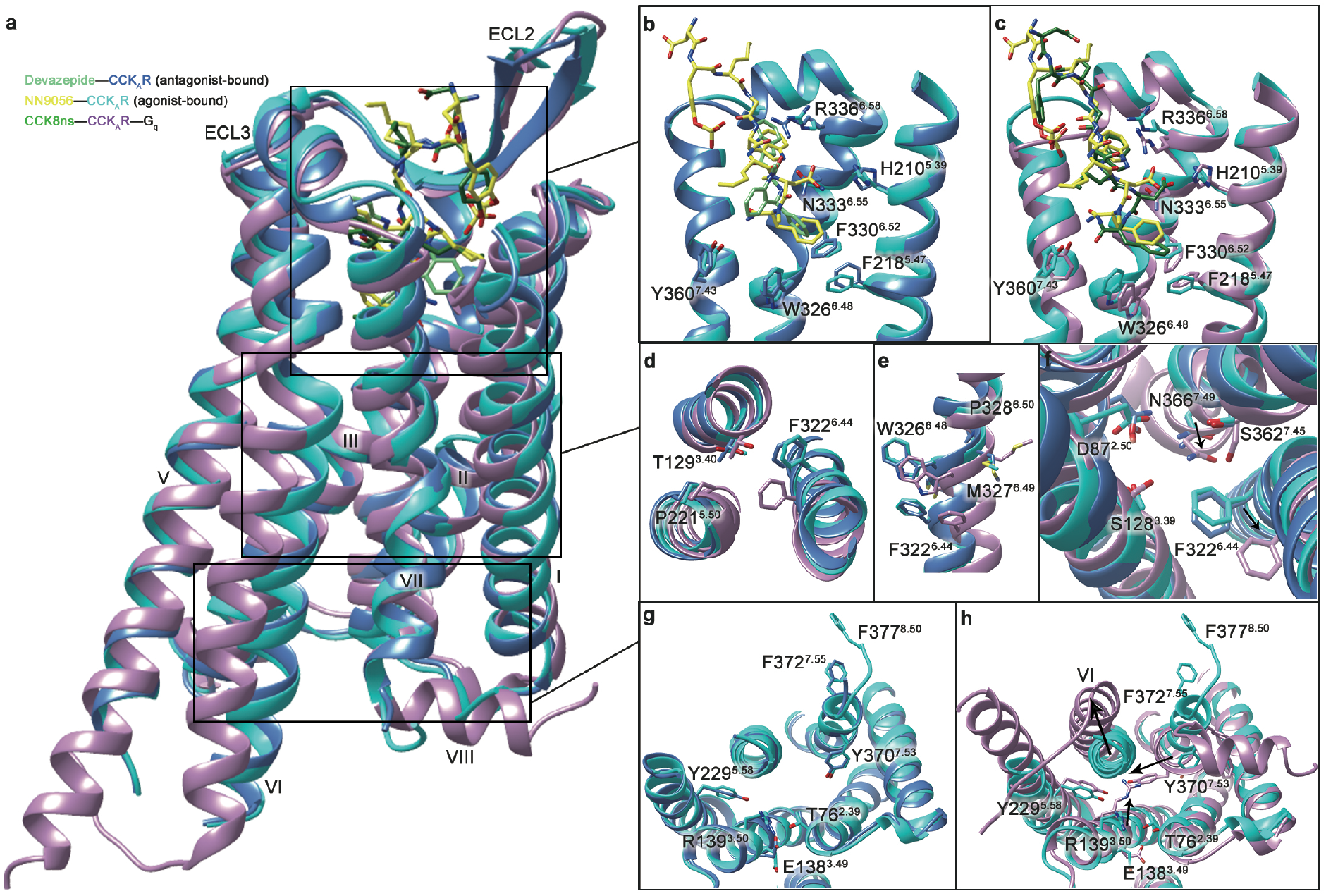
Activation process of CCK_A_R. **a**, Superimposition of the agonist-bound CCK_A_R structure with that bound to antagonist or both agonist and G_q_ (obtained in the accompanied paper). G protein is omitted for clarity. **b**, **c**, Conformational difference of the ligand-binding pocket among distinct states. **d-f**, Structural rearrangements of residue contacts occurred in PIF (**d**), FxxCWxP (**e**) and sodium pocket (**f**). **g**, **h**, Structural changes in the intracellular ends of helices VI and VII induced by agonist binding and G_q_ coupling. Conformational changes are indicated with black arrows.

The CCK_A_R–devazepide, CCK_A_R–NN9056 and the simulated CCK-8ns–CCK_A_R–G protein structures provide a systematic view of CCK_A_R in antagonist bound, agonist bound and agonist–G-protein bound states, which allows an in-depth analysis of the activation process of CCK_A_R. Structural comparison of CCK_A_R in complex with different types of ligands indicates that the position of the indole group in the ligands, which forms extensive interactions with ECL3, might play a key role in regulating receptor activity. The indole groups of devazepide and lintitript that mimic the side chain of the peptide residue W4 are 1.1 Å away from ECL3 compared to that in the peptide ligands (Fig. 3i). This difference suggests that conformational change in ECL3 may be involved in modulating receptor activation (Extended Data Fig. 6). Indeed, the extracellular half of the CCK_A_R structures of different active states overlap with each other except that ECL3 bends down by 2.9 Å (measured at the Cα carbon of G349), leading to a counterclockwise twisting of the extracellular regions of helices VI and VII (extracellular view) and side-chain reorientations of several key residues within the ligand-binding pocket (Fig. 4a–c). The inward movement of ECL3 allows the agonist to penetrate deeper in the binding pocket, forming a spatial hindrance and pushing the side chain of F330^6.52^. The downward movement of F330^6.52^ in turn influences the side chain of W326^6.48^ and induces an outward movement of the side chains of F218^5.47^, F322^6.44^ and F323^6.45^, thereby resulting in a significant increase of the peptide-receptor interface area (from 1,797 Å^2^ to 2,424 Å^2^). Besides W326^6.48^ and F322^6.44^, which were previously indicated as ‘transmission switch’ in FPR2^29^, the P230^5.50^ in the PIF motif also showed an inward shift upon activation. Consistently, several newly formed polar contacts including three hydrogen bonds between the CCK residue D7 and the residues H210^5.39^, N333^6.55^ and R336^6.58^ of the receptor were observed in the active CCK_A_R structure. Accompanying these movements initiated at the bottom of the ligand-binding pocket, both the sodium pocket and transmission switch (P^5.50^T^3.40^F^6.44^ and CWxP^6.50^ motifs) rearrange their residue contacts (Fig. 4d, e). By pointing to helix VI, S362^7.45^ forms two hydrogen bonds with W326^6.48^ and N366^7.46^, yielding a 2.4 Å-downward movement of W326^6.48^ and a 2.3 Å-inward movement of N366^7.46^ (measured at the Cα carbon of N366^7.46^), relative to those in the NN9056-bound structure. Despite no obvious sodium density was observed in all our CCKR structures, the distance between residues in sodium binding pocket such as D100^2.50^ and N366^7.49^ was closer upon activation (3.4 Å to 2.7 Å), forming a stronger interaction between helices II and VII. Meanwhile, the side chain of F322^6.44^ stretches out from the transmembrane domain core and the hydrophobic lock (L132^3.43^, I318^6.40^ and V319^6.41^) is broken, which further loosens the helices III-VI packing and facilitates the outward movement of the cytoplasmic end of helix VI.

On the intracellular side of the G protein-coupled CCK_A_R structure, helix VI of the receptor moves outwards by 6.0 Å (measured at the Cα carbon of A302^6.24^), while helix VI extends by ten residues and moves outwards by about 2.0 Å (Y237^5.66^ as a reference). In addition, helix VII rotates inward and moves toward helix III by 5.2 Å (measured at the Cα carbon of Y370^7.53^). In contrast to the restrains of the hydrogen bond with T76^2.39^ and the salt bridge with E138^3.49^ in the inactive state, R139^3.50^ exchanges these polar contacts upon agonist binding with interaction at the C terminus of α5 helix in the Gα subunit. Surprisingly, helix VIII of CCK_A_R was found in a non-canonical position that is perpendicular to the helical bundle without contacting any transmembrane helices in both agonist- or antagonist-bound structures. Upon G protein binding, helix VIII rotates toward helix I and resembles that of other class A GPCRs. Collectively, these movements create an intracellular crevice for G protein coupling.

In summary, we solved the CCK_A_R structures at different activation states and the structures of CCK_B_R in complex with different G proteins. Supported by receptor binding and signaling profile data, these structures help us better understand multiple aspects of CCK receptor biology including recognition of different types of ligands and peptide selectivity in the CCKR family, as well as the activation process of CCK_A_R.

## Methods

### Construct cloning

For crystallization studies of CCK_A_R, the human CCK_A_R gene was cloned into a modified pFastBac1 vector (Invitrogen), which contains an expression cassette with a hemagglutinin (HA) signal sequence followed by a Flag tag at the N terminus as well as a PreScission protease site followed by a 10 × His tag at the C terminus. To facilitate crystallization, T4L was inserted into between I240 and A302 at ICL3 of CCK_A_R with the truncation of the C-terminal residues G407–Q428. One mutation F130^3.41^W was introduced to solve the CCK_A_R–NN9056 complex structure. The construct was further optimized with D87^2.50^N to solve the structure of CCK_A_R–devazepide and CCK_A_R–lintitript complex structures.

For cryo-EM studies of CCK_B_R, the human CCK_B_R gene was cloned into a modified pFastBac1 vector (Invitrogen) with N-terminal HA signal sequence connected to a Flag-tag and C-terminal PreScission protease site followed by a 2 × Strep-tag. The construct was further optimized by truncation of C-terminal residues 419–447 to improve the protein yield and homogeneity. According to previous studies, a chimera G_q_ was generated by replacing N-terminal 30 amino acids with the N-terminal 24 amino acids of G_i1_^42^. A dominant-negative G_i2_ (DNG_i2_) gene was generated as previously described by introducing four Gα_i2_ subunit mutations, S47N, G204A, E246A and A327S. Both G_q_ and G_i2_ genes are cloned into the pFastBac1 vector. The human Gβ_1_ and Gγ_2_ subunits were integrated into the pFastBac Dual vector (Invitrogen) with an N-terminal 6 × His-tag at the N terminus of Gβ_1_. The scFv16 gene was cloned into a modified expression vector pFastBac1 with an N-terminal GP67 signaling peptide and a C-terminal 8 × His tag.

### Protein expression

The Bac-to-Bac baculovirus expression system (Invitrogen) was used to generate high-titer recombinant baculovirus (>10^9^ viral particles per ml). Expression of CCK_A_R was carried out by infection of *Spodoptera frugiperda* (*Sf*9) cells at a cell density of 2–3 × 10^6^ cells/ml with virus at a multiplicity of infection (MOI) of 5. Transfected cells were cultured at 27 °C for 48 h and then collected by centrifugation and stored at −80 °C until use.

For the CCK_B_R–gastrin-17–G_q_ complex, the modified CCK_B_R, Gα_q_, Gβ_1_γ_2_ and Ric8A (resistance to inhibitors of cholinesterase 8A), were co-expressed in High Five insect cells (Invitrogen) using the Bac-to-Bac baculovirus expression system. Cells at a density of 1.5 × 10^6^ cells per ml were infected with high-titer virus at a MOI ratio of 2:1:1:1 for CCK_B_R, Gα_q_, Gβ_1_γ_2_ and Ric8A. For the CCK_B_R–gastrin-17–G_i2_ complex, the modified CCK_B_R, Gα_i2_ and Gβ_1_γ_2_ were co-expressed in High Five cells using the Bac-to-Bac baculovirus expression system. Cells at a density of 3 × 10^6^ cells per ml were infected with high-titer virus at a MOI ratio of 2:1:1 for CCK_B_R, Gα_i2_ and Gβ_1_γ_2_. Cells were cultured at 27 °C and harvested 48 h after infection by centrifugation and stored at −80 °C until use.

### Purification of CCK_A_R-devazepide, CCK_A_R-lintitript and CCK_A_R-NN9056 complexes

Insect cells were disrupted by thawing frozen pellets in a hypotonic buffer containing 10 mM HEPES, pH 7.5, 10 mM MgCl_2_, 20 mM KCl and EDTA-free protease inhibitor cocktail (Roche) with the ratio of 1 tablet per 100 ml lysis buffer. Extensive washing was performed by repeated centrifugation in the same buffer and then in a high salt buffer containing 10 mM HEPES, pH 7.5, 10 mM MgCl_2_, 20 mM KCl and 1 M NaCl (two times each).

The purified membranes were thawed on ice in the presence of 2 mg/ml iodoacetamide and EDTA-free protease inhibitor cocktail, and incubated at 4 °C for 1 h before solubilization. The CCK_A_R–T4L was solubilized in a buffer containing 50 mM HEPES, pH 7.5, 300 mM NaCl, 0.5% (w/v) DDM, 0.1% (w/v) CHS, 10% glycerol with 50 μM lintitript, 40 μM devazepide, and 50 μM NN9056, respectively, at 4 °C for 3 h. The supernatant was isolated by centrifugation at 160,000 *g* for 30 min and incubated with TALON superflow metal affinity resin (Clontech) at 4 °C overnight. The resin was washed with 20 column volumes of wash buffer containing 25 mM HEPES, pH 7.5, 300 mM NaCl, 10% (v/v) glycerol, 0.05% (w/v) DDM, 0.01% (w/v) CHS, 30 mM imidazole and 50 μM lintitript, 40 μM devazepide, or 50 μM NN9056. The protein was eluted with 5 column volumes of elute buffer containing 50 mM HEPES, pH 7.5, 300 mM NaCl, 10% (v/v) glycerol, 0.05% (w/v) DDM, 0.01% (w/v) CHS, 300 mM imidazole and 50 μM lintitript, 40 μM devazepide, or 50 μM NN9056. A PD MiniTrap G-25 column (GE Healthcare) was used to remove imidazole. The protein was then treated overnight with His-tagged PreScission protease and His-tagged PNGase F to remove the C-terminal His tag and de-glycosylate the receptor. PreScission protease, PNGase F and the cleaved 10 × His tag were removed by passing the sample through Ni-NTA superflow resin (QIAGEN). The receptor was concentrated to 20–30 mg/ml with a 100 kDa cut-off concentrator (Millipore). Protein purity and mono-dispersity were examined by Nu-PAGE and analytical size-exclusion chromatography.

### Crystallization of CCK_A_R-devazepide, CCK_A_R-lintitript and CCK_A_R-NN9056 complexes

The CCK_A_R complex samples were crystallized using the lipid cubic phase (LCP) method by mixing 40% protein with 60% lipid (monoolein and cholesterol 10:1 by mass) using a syringe lipid mixer. After a clear LCP formed, the mixture was dispensed onto glass sandwich plates (Shanghai FAstal BioTech) into 40 nl drop and overlaid with 800 nl precipitant solution using a Gryphon robot (Art-Robbins). Crystals of the CCK_A_R–lintitript complex appeared after 2 days and grew to full size within 2 weeks in 0.1 M HEPES, pH 7.0-7.5, 20%–30% (v/v) PEG400, 200–300 mM sodium tartrate and 1%–2% 1,2-butanediol. The CCK_A_R–devazepide complex was crystallized in 0.1 M HEPES, pH 7.0-7.5, 20%–30% (v/v) PEG400 and 300–400 mM ammonium acetate. The CCK_A_R–NN9056 complex was crystallized in 0.1M HEPES, pH 7.0–7.5, 8%–12% (v/v) PPG400 and 50–100 mM ammonium acetate. Crystals were harvested from LCP using 50 μm micromounts (M2-L19-100/150, MiTeGen) and flash frozen in liquid nitrogen.

### Data collection and structure determination

X-ray diffraction data were collected at the SPring-8 beam line 41XU, Hyogo, Japan, using a 10 μm beam (at a wavelength of 1.0000 Å) and a Pilatus 3 6M detector. Crystals were exposed with a 10 μm × 8 μm beam for 0.2 s and 0.2 ° oscillation per frame. Data from the 34 best diffracting crystals of the CCK_A_R–lintitript complex, 17 crystals of the CCK_A_R–devazepide complex and 44 crystals of the CCK_A_R–NN9056 complex were processed using XDS^43^. Initial phase information was obtained by molecular replacement using the structures of NPY1R (PDB accession number 5ZBQ) and T4L (PDB accession number 1C6P), respectively, with the program Phaser^44^. All refinements were performed with Phenix^45^, followed by manual examination and rebuilding of the refined coordinates in the program COOT^46^ using both |2*Fo*| - |*Fc*| and |*Fo*| - |*Fc*| maps. The Ramachandran plot indicates that 94.5% (5.5%) of residues in the CCK_A_R–devazepide, 94.7% (5.3%) of residues in the CCK_A_R–lintitript and 90.3% (9.7%) of residues in the CCK_A_R-NN9056 complexes were in favored (allowed) region (no outliers). The final model of the CCK_A_R–devazepide complex contains 278 residues of CCK_A_R (K37–I240, A302–K375) and 160 residues (1–160) of T4 lysozyme; the final model of the CCK_A_R–lintitript complex contains 279 residues of CKK_A_R (K37–I240, A302–R376) and 160 residues (1–160) of T4 lysozyme; and the final model of the CCK_A_R-NN9056 complex contains 283 residues of CKK_A_R (K37–I240, A302–G380) and 160 residues (1–160) of T4 lysozyme. The remaining N- and C-terminal residues of CCK_A_R are disordered and were not modeled. Data collection and structure refinement results are shown in Extended Data Table 1.

### CCK_B_R–gastrin-17–G_q_ and CCK_B_R–gastrin-17–G_i2_ complex formation and purification

Cells were thawed on ice and suspended in a buffer containing 20 mM HEPES, pH 7.5, 50 mM NaCl, 2 mM MgCl_2_ and 100 μg/ml protease inhibitor (PanReac Applichem). Both CCK_B_R–gastrin-17–G_q_ and CCK_B_R–gastrin-17–G_i2_ complexes were formed on membrane by adding 50 μM gastrin-17 and 25 mU/ml apyrase (New England Bio-Labs), followed by incubation at 20 °C for 1 h. The membrane pellets were collected by ultra-centrifugation at 100,000 *g* for 30 min. The CCK_B_R–gastrin-17–G_q_ complex was extracted from the membrane using a buffer containing 50 mM HEPES, pH 7.5, 300 mM NaCl, 2 mM MgCl_2_, 0.5% (w/v) *n*-dodecyl-β-D-maltopyranoside (DDM, Anatrace), 0.1% (w/v) cholesteryl hemisuccinate (CHS, Sigma), 50 μM gastrin-17 and 25 mU/ml apyrase, while the CCK_B_R–gastrin-17–G_i2_ complex was extracted from the membrane using a buffer containing 50 mM HEPES, pH 7.5, 300 mM NaCl, 2 mM MgCl_2_, 0.5% (w/v) lauryl maltoseneopentyl glycol (LMNG, Anatrace), 0.05% (w/v) CHS, 50 μM gastrin-17 and 25 mU/ml apyrase. Both complexes were incubated at 4 °C for 3 h and the supernatants isolated by ultra-centrifugation at 100,000 *g* for 30 min, followed by incubation with pre-equilibrated Strep-Tactin Sepharose (IBA Lifesciences) at 4 °C overnight.

The resin with the immobilized CCK_B_R–gastrin-17–G_q_ complex was washed with 20 column volumes of washing buffer containing 25 mM HEPES, pH 7.5, 150 mM NaCl, 0.05% (w/v) DDM, 0.01% (w/v) CHS and 50 μM gastrin-17, while the resin with the immobilized CCK_B_R–gastrin-17–G_i2_ complex was washed with 20 column volumes of washing buffer containing 25 mM HEPES, pH 7.5, 150 mM NaCl, 0.01% (w/v) LMNG, 0.001% (w/v) CHS and 50 μM gastrin-17. Then, both complexes were subjected to the same purification protocol. The detergent was exchanged on resin with 10 column volumes of a buffer containing 25 mM HEPES, pH 7.5, 150 mM NaCl, 0.25% (w/v) glyco-diosgenin (GDN, Anatrace) and 50 μM gastrin-17 at 4 °C for 2 h. The resin was further washed with 20 column volumes of a buffer containing 25 mM HEPES, pH 7.5, 150 mM NaCl, 0.01% (w/v) GDN and 50 μM gastrin-17. The protein was then eluted with 5 column volumes of elute buffer containing 200 mM Tris-HCl, 500 mM NaCl, 0.01% GDN, 50 mM biotin and 50 μM gastrin-17. Purified complex was stabilized with addition of a 1.5 molar excess of scFv16 (preparation protocol shown as below) and incubated at 4 °C for 2 h. The complex protein was further purified by size exclusion chromatography on a Superdex 200 Increase 10/300 column (GE Healthcare) pre-equilibrated with 20 mM HEPES, pH 7.5, 150 mM NaCl, 0.01% (w/v) GDN (to remove uncoupled receptor), excess gastrin-17 and scFv16. The monomeric complex peak was collected and concentrated to ~5 mg/ml with a 100 kDa cut-off concentrator (Millipore), and then analyzed by SDS-PAGE and analytical size-exclusion chromatography.

### Expression and purification of scFv16

ScFv16 was expressed in High Five cells as a secreted protein using the baculovirus infection system. The culture medium with secreted scFv16 was harvested 48 h after infection and the protein was purified by affinity chromatography and size exclusion chromatography as previously described ^43^. Briefly, the cell culture supernatant was pH balanced with addition of 20 mM Tris-HCl, pH 8.0 and the chelating agents was removed by adding 1 mM nickel chloride and 5 mM calcium chloride followed by incubation at 20 °C for 1 h. Precipitates were removed by ultra-centrifugation at 100,000 *g* for 30 min and the supernatant was incubated with TALON resin at 4 °C overnight. The resin was washed with 20 column volumes of washing buffer I containing 20 mM HEPES, pH 7.5, 500 mM NaCl and 10 mM imidazole, and then was further washed with 20 column volumes of washing buffer II containing 20 mM HEPES, pH 7.5, 100 mM NaCl and 10 mM imidazole. The protein was eluted in elution buffer containing 20 mM HEPES, pH 7.5, 100 mM NaCl and 250 mM imidazole. Imidazole was removed using PD MiniTrap G-25 column (GE Healthcare). The protein was then treated with His-tagged PreScission protease to remove the C-terminal 8 × His tag at 4 °C overnight. Cleaved protein was further purified by reloading into Ni-NTA resin (QIAGEN). The flow through was collected, concentrated to ~3 mg/ml with a 10 kDa cut-off concentrator (Millipore), flash-frozen by liquid nitrogen and stored at −80 °C until use.

### Cryo-EM data acquisition and processing

Negative staining EM was used to confirm the formation of the CCK_B_R–gastrin-17–G_q_/G_i2_ complexes. The protein quality of the complexes was evaluated by 200 kV cryo-EM. For 300 kV cryo-EM, 3 μl of protein sample (5 mg/ml) was applied to glow-discharged 300 mesh gold grids (CryoMatrix M024-Au300-R12/13) and vitrified using a FEI Vitrobot Mark IV (ThermoFisher Scientific), at 4 °C and 100% humidity with blot time of 1 s and blot force of 0 followed by flash-frozen in liquid ethane. Images were collected using a Titan Krios electron microscope operated at 300 kV with a K3 Summit direct electron detector (Gatan) at a nominal magnification of × 81,000, corresponding to a pixel size of 1.045 Å. The slit width for zero loss peak was 20 eV. Defocus values are ranged from −0.8 μm to −1.5 μm. Each movie comprises 40 frames in a total of 3 s with 0.075 s exposure per frame, and the total dose is 70 electrons per Å^2^. Automated single-particle data acquisition was performed with SerialEM^47^.

Collected images of the CCK_B_R–gastrin-17–G_q_/G_i2_ complex samples were subjected to beam-induced motion correction using MotionCor2. Contrast transfer function (CTF) parameters for each image were determined by Gctf v1.18^48^. Guided by a template generated from initial auto-picking, the particles were extracted by auto-picking using both RELION3.1^49^ and Gautomatch v0.56 (developed by K. Zhang, MRC Laboratory of Molecular Biology, Cambridge, UK, http://www.mrc-lmb.cam.ac.uk/kzhang/Gautomatch/). 2D classification, 3D classification, 3D auto-refinement, Bayesian polishing and CTF refinement were performed using RELION3.1. The resolutions of density maps were calculated by the gold-standard Fourier Shell Correlation (FSC) using the 0.143 criterion. Local resolution estimation was determined by ResMap v1.1.4.

For the CCK_B_R–gastrin-17–G_q_ complex, a total of 5,625 images were collected, followed by beam-induced motion correction and CTF determination. All 3,632,729 particles were subjected to two rounds of reference-free 2D classification to discard false positive particles. An ab initio model generated by RELION3.1 was used as initial reference model for 3D classification. A subset of 418,664 particles were selected for another round of 3D classification that focused the alignment on the complex. The best-looking dataset of 354,647 particles were subjected to 3D auto-refinement, resulting in an initial 3.7 Å density map. A final 3.1 Å map was sharpened by postprocess with a B-factor of –75 Å^2^.

For the CCK_B_R–gastrin-17–G_i2_ complex, two datasets (6,471 images and 5,072 images) were collected and each dataset was individually subjected to beam-induced motion correction, CTF determination, auto-picking, 2D classification, 3D classification and 3D auto-refinement, resulting in initial 4.1 Å and 4.0 Å density maps, respectively. After separate Bayesian polishing, the two subsets (1,067,650 particles and 270,503 particles) were combined and subjected to another round of 3D auto-refinement. CTF refinement and postprocess with a B-factor of –111 Å^2^ were performed subsequently and result in a final 3.3 Å map.

### Model building and refinement

The model of the CCK_B_R–gastrin-17–G_q_ complex was built using the receptor from the CCK_A_R–NN9056 crystal structure, the G_q_ from 25-CN-NBOH-HTR2A-mini G_q_ cryo-EM structure (PDB: 6WHA) and YM-254890–G_q_ crystal structure (PDB: 3AH8), and the Gβ, Gγ and scFv16 from glucagon–GCGR–G_i_ cryo-EM structure (PDB: 6LML) as initial models, respectively. The model of the CCK_B_R–gastrin-17–G_i2_ complex was built using the receptor from CCK_A_R–NN9056 crystal structure and the G protein heterotrimer from the glucagon– GCGR–G_i_ cryo-EM structure (PDB: 6LML) as initial models. Both models were docked into the corresponding cryo-EM density map using ChimeraX v1.1^50^, followed by iterative manual adjustment in COOT^46^ and real space refinement using phenix.real_space_refine in PHENIX^45^. The model statistics was validated using Molprobity. The final model of the CCK_B_R–gastrin-17–G_i2_ complex contains 275 residues of CCK_B_R (A55–L249, L326–C405), 217 residues of Gα_i2_ (K10–M53, T183–F355), 306 residues of Gβ_1_ (N35–N340), 33 residues of Gγ_2_ (K29–F61) and 17 residues of gastrin-17. For the CCK_B_R–gastrin-17–G_q_ complex, the final model contains 275 residues of CCK_B_R (A55–L249, L326–C405), 224 residues of Gα_q_ (A7–G58, G182–V353), 306 residues of Gβ_1_ (N35–N340), 33 residues of Gγ_2_ (K29–F61), 232 residues of antibody scFv16 (V19–S138, S153–L264), and 17 residues of gastrin-17. For both models, the remaining N- and C-terminal residues of CCK2 and heterotrimeric G_i_/G_q_ are disordered and were not modeled. The final refinement statistics are provided in Extended Data Table 2. The extent of any model overfitting during refinement was measured by refining the final model against one of the half-maps and by comparing the resulting map versus model FSC curves with the two half-maps and the final model.

### Radiolabeled ligand-binding assay

The wild-type (WT) or mutant CCKRs were transiently transfected into HEK 293T cells (purchased from the Cell Bank at the Chinese Academy of Sciences) which were cultured in poly-D-lysine coated 96-well plates. Twenty-four hours later, the cells were washed twice and incubated with blocking buffer (DMEM medium supplemented with 33 mM HEPES, and 0.1% (w/v) BSA, pH 7.4) for 2 h at 37 °C. After three times washes by cold-ice PBS, the cells were treated by a constant concentration of ^125^I-CCK-8 (30 pM, PerkinElmer) plus 8 different concentrations of gastrin-17 (128 pM–10 μM) for 3 h at room temperature (RT). Cells were washed three times with ice-cold PBS and lysed by 50 μl lysis buffer (PBS supplemented with 20 mM Tris-HCl and 1% (v/v) Triton X-100, pH 7.4). Subsequently, the plates were counted for radioactivity (counts per minute, CPM) in a scintillation counter (MicroBeta^2^ plate counter, PerkinElmer) using 150 μl scintillation cocktail (OptiPhase SuperMix, PerkinElmer).

### Inositol phosphate accumulation assay

Inositol phosphate accumulation was measured using an IP-One G_q_ assay kit (Cisbio Bioassays). The WT and mutant CCKRs were cloned into pTT5 vector (Invitrogen) and transiently transfected into HEK 293F cells. After 48 h expression, the cells were plated into 384-well plates (8,000 cells per well). For agonist effects, various gradient concentrations of NN9056 or CCK-8 (1 pM to 10 μM diluted in stimulation buffer) were added and incubated at 37 °C for 1 h. IP1-d2 and anti-IP1 cryptate were then applied and incubated at RT for 1 h. For antagonist experiments, 1 μM lintitript or 10 μM devazepide was introduced and incubated at 37 °C for 1 h followed by addition of different concentrations of NN9056 (10 pM to 100 μM diluted in stimulation buffer) and 1 h incubation at 37 °C. Fluorescence signals were read by Synergy H1 plate reader (Biotech) with excitation at 330 nm and emission at 620 and 665 nm. The inositol phosphate accumulation curves, EC50 and pEC50 ± S.E.M. were calculated using nonlinear regression curve fitting in Prism 8.

### Molecular dynamics (MD) simulation

To simulate the CCK_A_R in complex with CCK-8 in a no-fusion/no-mutation background, the crystal structure of CCK_A_R–NN9056 was prepared by the Protein Preparation Wizard (Schrodinger 2017-4). The initial conformation of CCK-8 with CCK_A_R was constructed on that of NN9056 by multiple rounds of single residue modification/mutation and energy minimization using the Protein Preparation Wizard. Residues D87^2.50^ and E138^3.49^ were protonated to simulate the protonation upon receptor activation while all other titratable residues were left in their dominant protonation state at pH 7.0. The complexes were embedded in a bilayer composed of 148 POPC lipids and solvated with 0.15 M NaCl in explicitly represented water using CHARMM-GUI Membrane Builder^51^. The CHARMM36-CAMP force filed was adopted for protein, peptides, lipids and salt ions, while the CHARMM TIP3P model was chosen for water. Parameters for the sulfated tyrosine was generated using the CHARMM General Force Field (CGenFF), program version 1.0.0. MD simulations were performed by Gromacs 2018.5. The Particle Mesh Ewald (PME) method was used to treat all electrostatic interactions beyond a cutoff of 10 Å and the bonds involving hydrogen atoms were constrained using LINCS algorithm. The constructed system was firstly relaxed using the steepest descent energy minimization, followed by slow heating of the system to 310 K with restraints. The restraints were reduced gradually over 18 ns, with a simulation step of 1 fs. Finally, the system was run without restraints, with a time step of 2 fs in the NPT ensemble at 310 K and 1 bar using the v-rescale thermostat and the semi-isotropic Parrinello-Rahman barostat, respectively. For each system, four independent 500 ns production simulations were performed, and trajectory were saved every 50 ps.

### Data availability

Atomic coordinates for the structures of CCK_A_R-lintitript, CCK_A_R-devazepide and CCK_A_R-NN9056 have been deposited in the RCSB PDB under accession codes ####, #### and ####. Atomic coordinates and cryo-EM density maps for the structures of inactive CCK_B_R-gastrin-G_i_ and CCK_B_R-gastrin-G_q_ have been deposited in the RCSB Protein Data Bank (PDB) under accession codes #### and ####, and the Electron Microscopy Data Bank (EMDB) under accession codes EMD-#### and EMD-####.

## Acknowledgements

This work was partially supported by the National Key R&D Programs of China 2018YFA0507000 (H.X.X., S.Z., M.-W.W., B.W. and Q.Zhao); the National Science Foundation of China grants 21704064 (Q.Z.), 81773792 and 81973373 (D.Y.), 31800621 (S.H.), 31770796 (Y.J.), 31971178 (S.Z.), 81872915 and 82073904 (M.-W.W.), 31825010 (B.W.) and 81525024 (Q.Zhao); National Science and Technology Major Project of China – Key New Drug Creation and Manufacturing Program 2018ZX09735–001 (M.-W.W.) and 2018ZX09711002–002–005 (D.Y.); and CAS Strategic Prority Research Program XDB37000000 (B.W.). The synchrotron radiation experiments were performed at the BL41XU of SPring-8 with approval of the Japan Synchrotron Radiation Research Institute (Proposal no. 2019A2543, 2019B2543, 2019A2541 and 2019B2541). We thank the beamline staff members K. Hasegawa, N. Mizuno, T. Kawamura, and H. Murakami of the BL41XU for help with X-ray data collection. The cryo-EM data were collected at the Cryo-Electron Microscopy Research Center, Shanghai Institute of Materia Medica (SIMM). The authors thank the staff at the SIMM Cryo-Electron Microscopy Research Center for their technical support.

## Author contributions

X.Z. optimized the construct, purified the CCK_A_R protein, performed crystallization trials, solved the structure, performed signaling assays, and helped manuscript preparation. C.H. optimized the construct, purified the CCK_B_R protein, performed crystallization trials, solved the structure, performed signaling assays, and helped manuscript preparation. M.W. processed the cryo-EM data and solved the structures of CCK_B_R. D.Y., W.F., A.D. and J.W. designed and performed the receptor bingding and functional assays. Q.Z. performed molecular dynamics simulation and docking studies, and helped manuscript preparation. Y.Z. helped signaling assay design and manuscript preparation. H.Z. collected X-ray diffraction data. X.C. helped protein expression. Z.Y. participated in manuscript preperation. Y.J. solved the cryo-EM structures of CCK_A_R–G protein complexes. U.S. designed and synthesized the ligands. Q.T. asissited in the cryo-EM data collection. S.H. helped structure determination. S.R. helped ligand selection and data analysis. H.E.X. helped analysis of cryo-EM structures of CCK_A_R–G protein complexes and manuscript preparation. S.Z. oversaw molecular dynamics simulation and docking studies, and edited the manuscript. M.-W.W. oversaw the binding and signaling assays, and edited the manuscript. B.W. and Q.Zhao initiated the project, planned and supervised the research, and wrote the manuscript with inputs from all co-authors.

## Extended Data Figures

**Extended Data Fig. 1.**
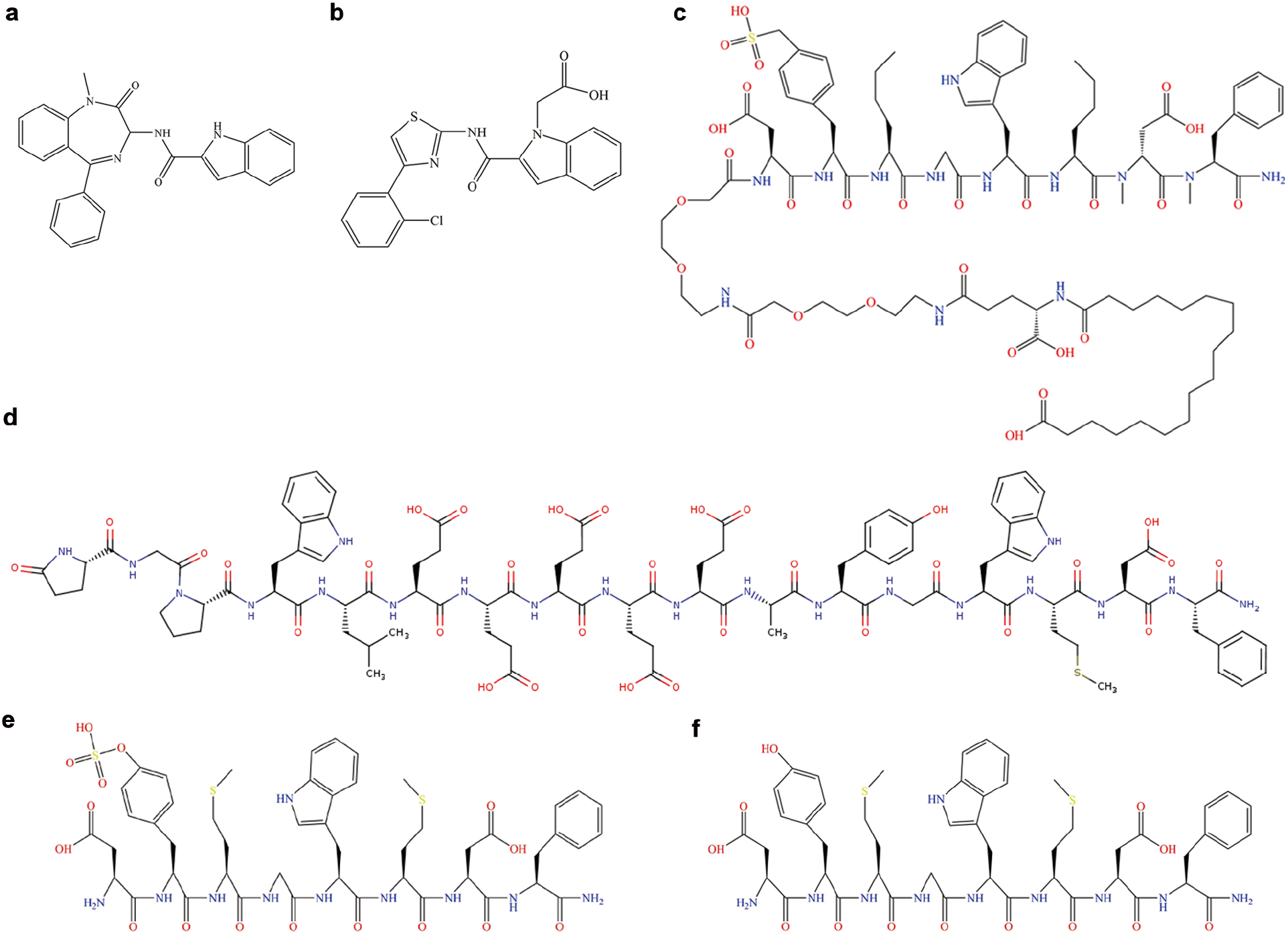
Structures of the ligands in the CCK_A_R and CCK_B_R complexes and molecular dynamic simulations. **a**, Devazepide. **b**, Lintitript. **c**, NN9056. **d**, Gastrin-17. **e**, CCK-8. **f**, CCK-8ns.

**Extended Data Fig. 2.**
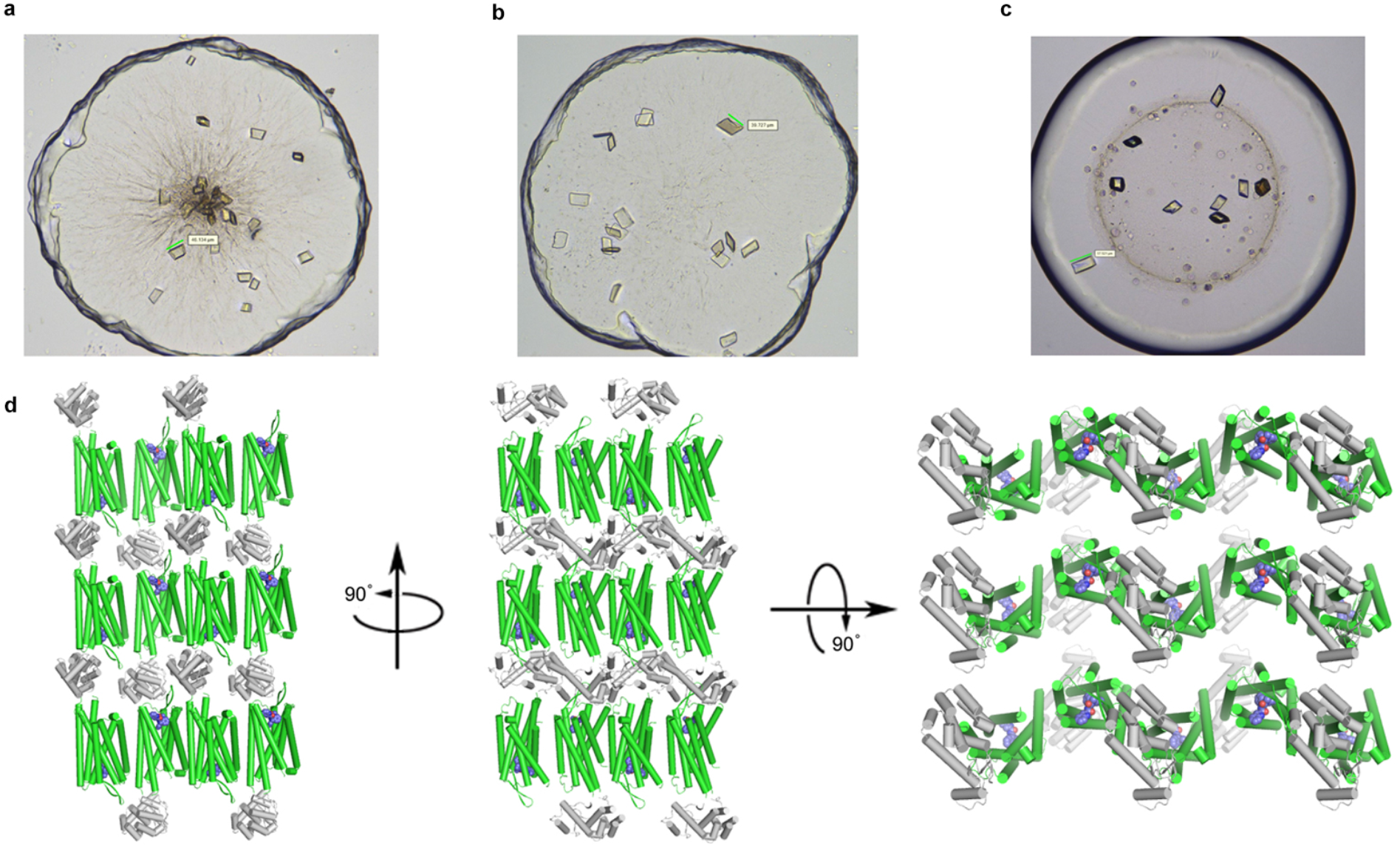
CCK_A_R crystals and lattice packing. **a**–**c**, Crystals of CCK_A_R–T4L in complex with devazepide (**a**), lintitript (**b**) and NN9056 (**c**). **d**, Lattice packing of CCK_A_R–T4L–devazepide crystals with CCK_A_R depicted in green, T4L in grey and devazepide shown as spheres in blue. The main contacts contained nonpolar interactions among CCK_A_R molecules mediated by helices I and V and interactions between ECL2 and ECL3 of CCK_A_R and T4L. The packing patterns of CCK_A_R–T4L–lintitript and CCK_A_R–T4L–NN9056 are same as CCK_A_R–T4L–devazepide.

**Extended Data Fig. 3.**
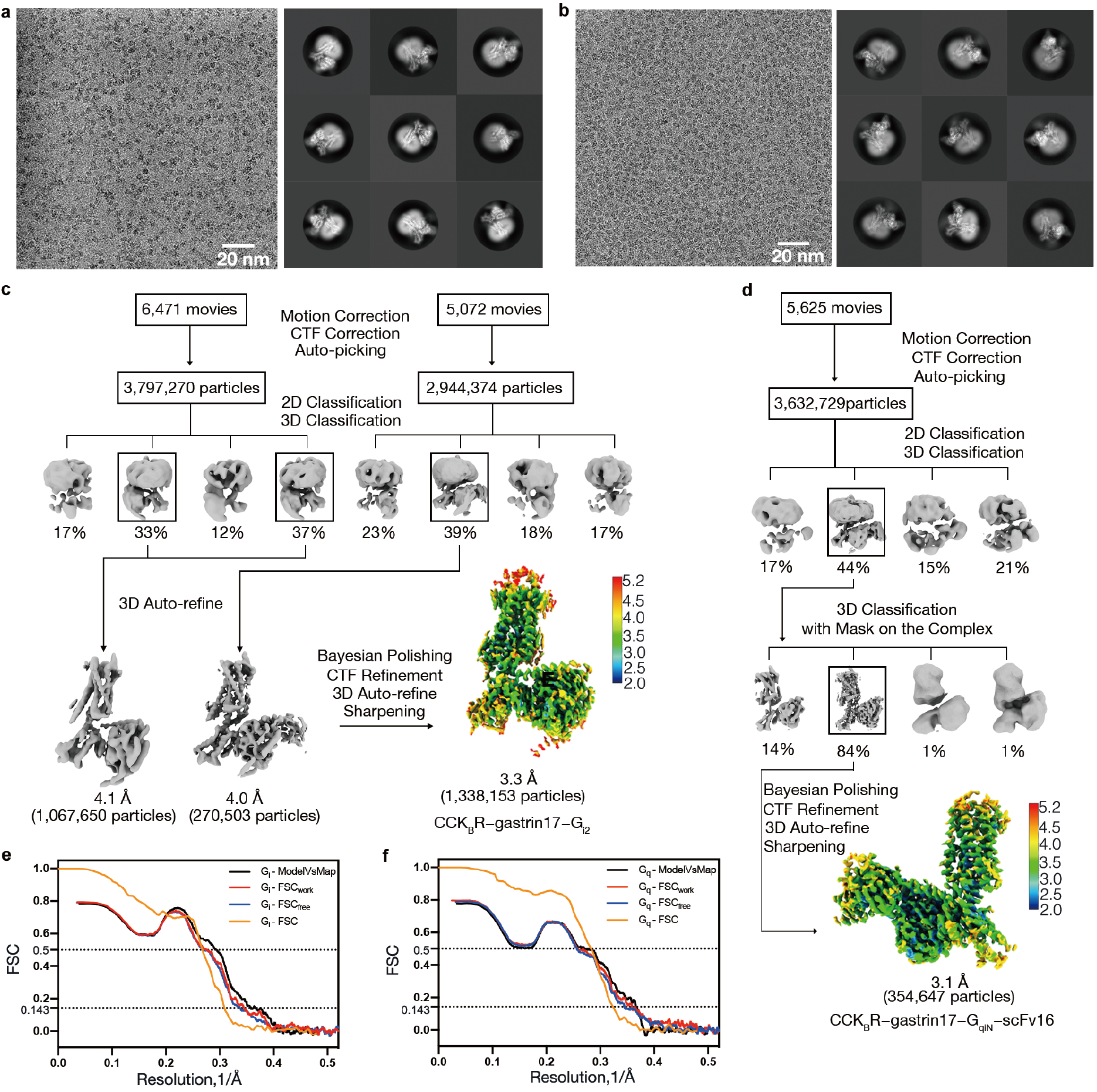
Cryo-EM data processing of the CCK_B_R–gastrin-17–G_i2_ and CCK_B_R–gastrin-17–G_q_ complexes. **a**, Representative cryo-EM image and 2D averages of the CCK_B_R–gastrin-17–G_i2_ complex. **b**, Representative cryo-EM image and 2D averages of CCK_B_R–gastrin-17–G_q_ complex. **c**, Workflow of cryo-EM data processing with cryo-EM map colored according to local resolution (Å) for the CCK_B_R–gastrin-17–G_i2_ complex. **d**, Workflow of cryo-EM data processing with cryo-EM map colored according to local resolution (Å) for the CCK_B_R–gastrin-17–G_q_ complex. **e**, Gold-standard FSC curve and cross-validation of model to cryo-EM density map of CCK_B_R–gastrin-17–G_i2_. FSC curves for the final model versus the final map, FSCwork curve, FSCfree curve and FSC curves are shown in black, red, blue and orange, respectively. **f**, Gold-standard FSC curve and cross-validation of model to cryo-EM density map of CCK_B_R–gastrin-17–G_qiN_.

**Extended Data Fig. 4.**
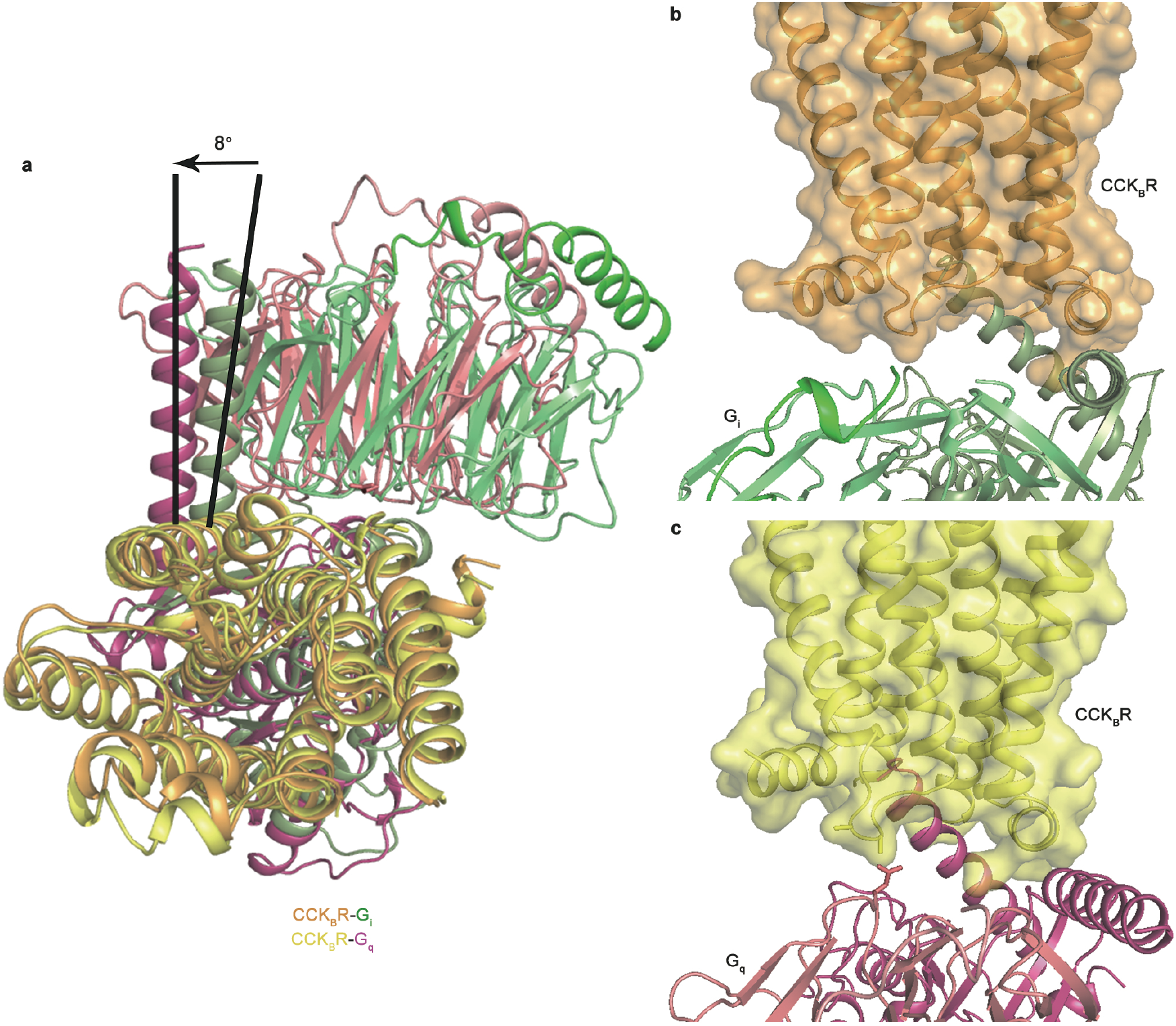
G_i2_ and G_q_ binding comparison in the CCK_B_R–gastrin-17 complexes. **a**, Superposition of the gastrin-17–CCK_B_R–G_i2_ and gastrin-17–CCK_B_R–G_q_ complex structures. The structures are aligned based on receptor region. The CCK_B_R are shown in wheat and light orange cartoons for G_i2_ and G_q_ complexes. The G_i2_ trimers are shown in blue, slate and light blue cartoons, and the G_q_ trimers are shown in yellow, light yellow and limon cartoons, respectively. Side view of G_i2_ (**b**) and G_q_ (**c**) binding to the receptor from the same angle. The structures are colored according to panel (**a**).

**Extended Data Fig. 5.**
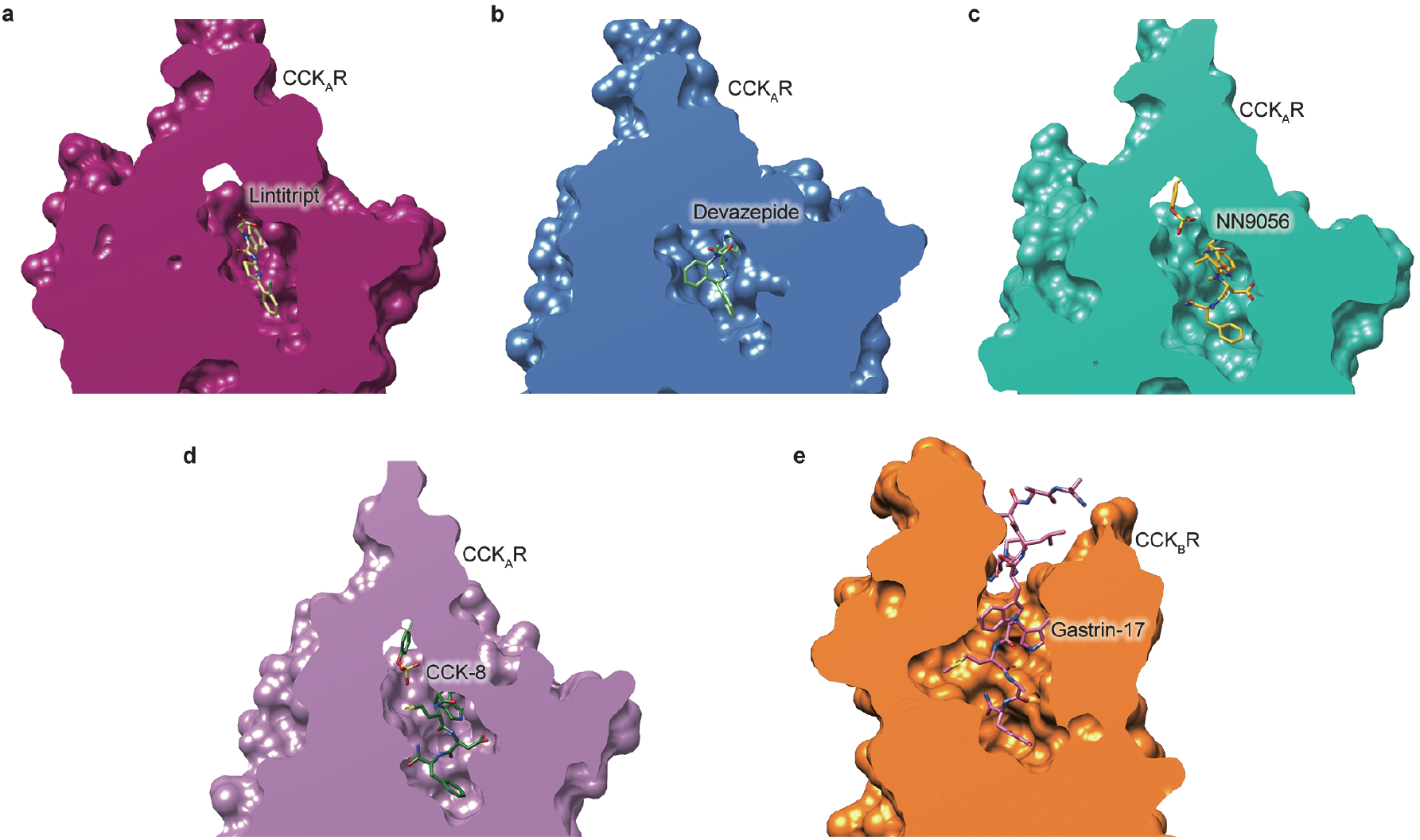
Cut-view of orthosteric sites of CCK_A_R and CCK_B_R. **a**, CCK_A_R–lintitript. **b**, CCK_A_R–devazepide. **c**, CCK_A_R–NN9056.**d**, CCK_A_R–CCK-8.**e**, CCK_B_R–gastrin-17.

**Extended Data Fig. 6.**
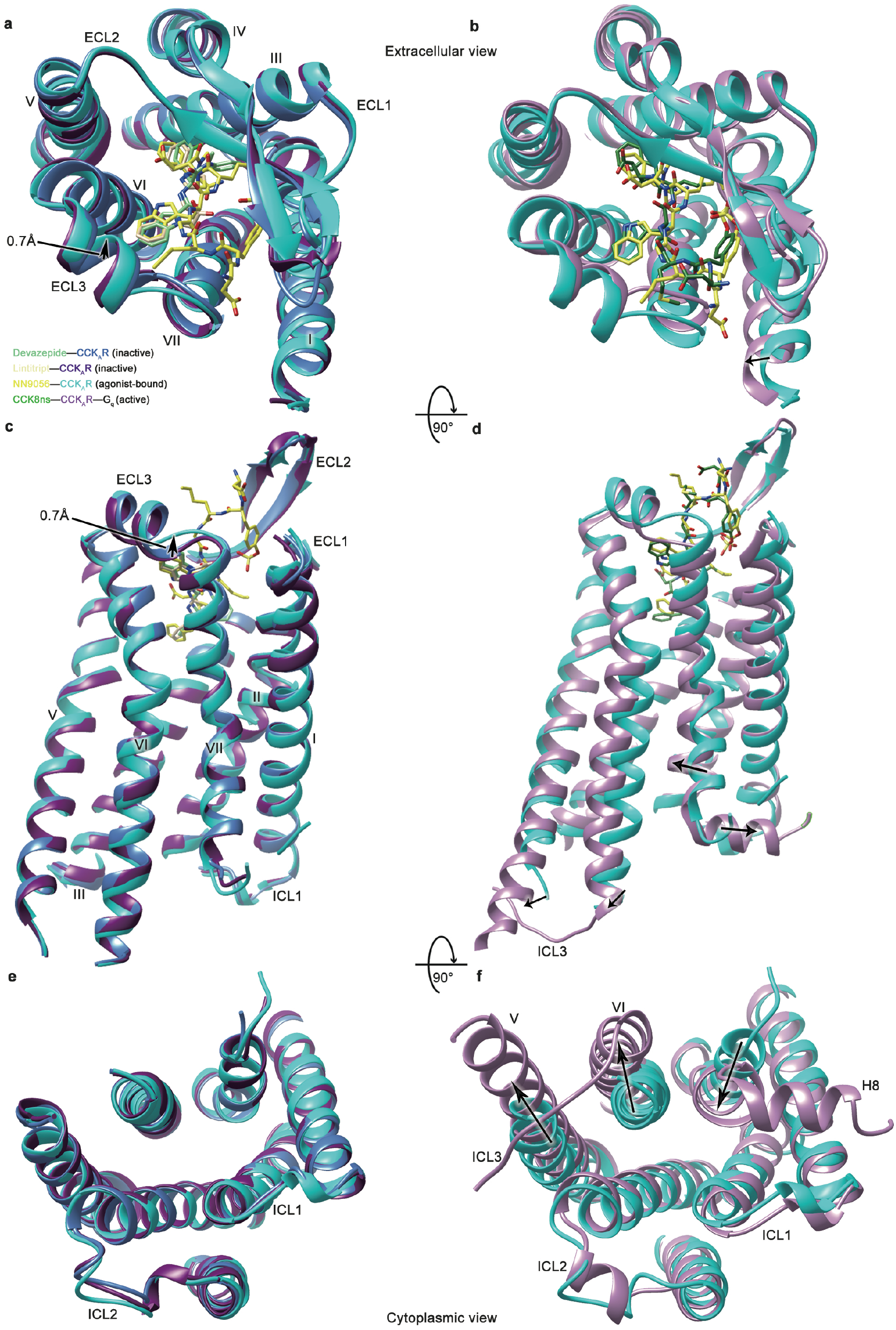
Comparison of the CCK_A_R structures. Extracellular (top), side (middle) and intracellular (bottom) views of agonist NN9056-bound (violet), antagonist-bound (lime green for devazepide or pale green for lintitript), and both agonist and G protein bound (orange) CCK_A_R structures. G proteins are omitted for clarity. Conformational changes are indicated with black arrows.

**Extended Data Table 1.** X-ray data collection and refinement statistics of the CCK_A_R–devazepide, CCK_A_R–lintitript and CCK_A_R–NN9056 complex structures.

|  | CCK<sub>A</sub>R–devazepide<sup>a</sup> | CCK<sub>A</sub>R–lontitript<sup>b</sup> | CCK<sub>A</sub>R–NN9056<sup>c</sup> |
| --- | --- | --- | --- |
| **Data collection** |  |  |  |
| Space group | *P2<sub>1</sub>* | *P2<sub>1</sub>* | *P2<sub>1</sub>* |
| Cell dimensions |  |  |  |
| a, b, c (Å) | 54.8, 72.4, 86.1 | 54.4, 72.0, 86.0 | 56.6, 72.5, 87.9 |
| α, β, γ (°) | 90.0, 107.3, 90.0 | 90.0, 106.1, 90.0 | 90.0, 100.5, 90.0 |
| Resolution (Å) | 29.8-2.5 (2.64-2.50) | 29.6-2.8 (2.95-2.80) | 29.5-3.0 (3.16-3.00) |
| *R<sub>merge</sub>* (%) | 18.1 (89.2) | 19.0 (70.3) | 20.6 (57.3) |
| Mean *I*/σ*I* | 5.0 (1.1) | 5.2 (1.1) | 4.5 (1.0) |
| Completeness (%) | 98.2 (99.1) | 97.5 (96.7) | 97.1 (96.7) |
| Redundancy | 4.8 (4.6) | 4.1 (3.9) | 4.4 (4.2) |
| **Refinement** |  |  |  |
| Resolution (Å) | 29.8-2.5 | 29.6-2.8 | 29.5-3.0 |
| No. reflections | 22,038 (1,618) | 15,523 (1,158) | 26,470 (1,931) |
| *R<sub>work</sub>*/*R<sub>free</sub>* (%) | 0.209 / 0.264 | 0.234 / 0.250 | 0.216 / 0.265 |
| No. atoms |  |  |  |
| Protein | 3,439 | 3,453 | 3,488 |
| Ligand/ion | 31 | 28 | 80 |
| Average B-factors (Å<sup>2</sup>) |  |  |  |
| Protein | 81.6 | 91.3 | 113.9 |
| Ligand | 69.2 | 79.7 | 102 |
| R.m.s. deviations |  |  |  |
| Bond lengths (Å) | 0.008 | 0.010 | 0.008 |
| Bond angles (°) | 1.09 | 1.29 | 1.46 |

<sup>a</sup>17 crystals were used for structure determination. Values in parentheses are for highest-resolution shell. <sup>b</sup>34 crystals were used for structure determination. Values in parentheses are for highest-resolution shell. <sup>c</sup>44 crystals were used for structure determination. Values in parentheses are for highest-resolution shell.

**Extended Data Table 2.** Cryo-EM data collection and refinement statistics of the gastrin-17–CCK_B_R–G_i2_ and gastrin-17–CCK_B_R–G_q_ complex structures.

|  | CCK<sub>B</sub>R–gastrin-17–G<sub>i2</sub> | CCK<sub>B</sub>R–gastrin-17–G<sub>q</sub> |
| --- | --- | --- |
| **Data collection and processing** |  |  |
| Magnification | 81,000 | 81,000 |
| Voltage (kV) | 300 | 300 |
| Electron exposure (e<sup>-</sup>/Å<sup>2</sup>) | 70 | 70 |
| Defocus range (μm) | -0.8 ~ -1.5 | -0.8 ~ -1.5 |
| Pixel size (Å) | 1.045 | 1.045 |
| Symmetry imposed | C1 | C1 |
| Initial particle projections (no.) | 6,741,644 | 3,632,729 |
| Final particle projections (no.) | 1,338,153 | 354,647 |
| Map resolution (Å) | 3.3 | 3.1 |
| FSC threshold | 0.143 | 0.143 |
| Map resolution range (Å) | 2.5–5.0 | 2.5–5.0 |
| **Refinement** |  |  |
| Initial model used (PDB code) | CCK<sub>A</sub>R–NN9056* | CCK<sub>A</sub>R–NN9056* |
|  | 6LML | 6WHA,3AH8,6LML |
| Model resolution (Å) | 3.4 | 3.6 |
| FSC threshold | 0.5 | 0.5 |
| Map sharpening B factor (Å<sup>2</sup>) | –111 | –75 |
| **Model composition** |  |  |
| Non-hydrogen atoms | 6601 | 8486 |
| Protein residues | 848 (6601 atoms) | 1087 (8486 atoms) |
| Receptor residues | 275 (2128 atoms) | 275 (2128 atoms) |
| G protein residues | 556 (4340 atoms) | 563 (4442 atoms) |
| Antibody residues | -- | 232 (1783 atoms) |
| Ligand residues | 17 (133 atoms) | 17 (133 atoms) |
| **B factors (Å<sup>2</sup>)** |  |  |
| Protein | 57.08 | 62.02 |
| Ligand | 87.27 | 109.52 |
| **R.m.s. deviations** |  |  |
| Bond lengths (Å) | 0.006 | 0.009 |
| Bond angles (°) | 0.986 | 1.589 |
| **Validation** |  |  |
| MolProbity score | 1.63 | 1.74 |
| Clashscore | 7.36 | 8.93 |
| Rotamer outliers (%) | 0.00 | 0.00 |
| **Ramachandran plot** |  |  |
| Favored (%) | 96.52 | 96.16 |
| Allowed (%) | 3.48 | 3.84 |
| Disallowed (%) | 0.00 | 0.00 |

**Extended Data Table 3.** Effects of devazepide and lintitript on IP1 accumulation in wild-type and mutant CCK_A_Rs.

| Mutant | NN9056 | NN9056/devazepide<sup>a</sup> |  | NN9056/lintitript<sup>a</sup> |  | Expression<sup>c</sup><br>(% of WT) |
| --- | --- | --- | --- | --- | --- | --- |
|  | EC<sub>50</sub> (nM) | EC<sub>50</sub> (nM) | Ratio<sup>b</sup> | EC<sub>50</sub> (nM) | Ratio<sup>b</sup> |  |
| Wild-type | 13 | 1,235 | 98 | 770 | 61 | 100 |
| N98<sup>2.61</sup>A | 28 | 10,060 | 359 | /<sup>d</sup> | / | 222 ± 8 |
| T117<sup>3.28</sup>A | 24 | 982 | 41 | /<sup>d</sup> | / | 97 ± 6 |
| M121<sup>3.32</sup>A | 28 | 2,219 | 79 | 577 | 21 | 149 ± 8 |
| Y176<sup>4.60</sup>A | 5.3 | 1,942 | 366 | 602 | 116 | 41 ± 3 |
| F330<sup>6.52</sup>A | 5.4 | 198 | 37 | 393 | 73 | 43 ± 4 |
| N333<sup>6.55</sup>A | 8.7 | 324 | 37 | 84 | 10 | 123 ± 20 |
| R336<sup>6.58</sup>A | 19 | 710 | 37 | 151 | 8 | 139 ± 7 |
| L347<sup>7.30</sup>A | 26 | 2,090 | 80 | 280 | 11 | 49 ± 8 |
| I352<sup>7.35</sup>A | 10 | 816 | 82 | 289 | 29 | 134 ± 18 |

<sup>a</sup>EC<sub>50</sub> values were determined after 1 h stimulation with increasing concentrations of NN9056 together with 10 μM devazepide or 1 μM lintitript. All values are performed at least three independent experiments performed in triplicate.
<sup>b</sup>The EC<sub>50</sub> ratio represents the shift between NN9056 and NN9056 plus antagonist curve ( $EC_{50(NN9056+antagonist)}/EC_{50(NN9056)}$ ) and characterizes the antagonistic effect. A reduced EC<sub>50</sub> ratio of mutant compared to the wild-type receptor was interpreted as important of the respective antagonist.
<sup>c</sup>Protein expression levels of CCK<sub>A</sub>R constructs at the cell surface were determined in parallel by flow cytometry with an anti-FLAG antibody and reported as percent compared to the WT CCK<sub>A</sub>R from at least three independent measurements performed in duplicate.
<sup>d</sup>The EC<sub>50</sub> was not measured due to lack of interaction in the structure.

**Extended Data Table 4.** Binding of devazepide and lintitript to wild-type (WT) and mutant CCK_A_Rs in competition with ^125^I-CCK-8.

| Mutant | Devazepide |  |  | Lintitript |  |  | N<sup>c</sup> |
| --- | --- | --- | --- | --- | --- | --- | --- |
|  | IC<sub>50</sub><br>(nM) | pIC<sub>50</sub> ±<br>S.E.M.<sup>a</sup> | Span<br>(% WT)<sup>b</sup> | IC<sub>50</sub><br>(nM) | pIC<sub>50</sub> ±<br>S.E.M. | Span<br>(% WT) |  |
| Wild-type | 17.0 | 7.77 ± 0.09 | 100 | 14.2 | 7.85 ± 0.07 | 100 | 3 |
| N98<sup>2.61</sup>A | 17.0 | 7.77 ± 0.08 | 66 ± 2 | /<sup>d</sup> | / | / | 3 |
| T117<sup>3.28</sup>A | 18.6 | 7.73 ± 0.09 | 147 ± 6 | / | / | / | 3 |
| T118<sup>3.29</sup>A | 13.1 | 7.88 ± 0.07 | 105 ± 3 | / | / | / | 3 |
| M121<sup>3.32</sup>A | 18.6 | 7.73 ± 0.10 | 175 ± 8 | 190.0 | 6.72 ± 0.07 | 160 ± 6 | 3 |
| Y176<sup>4.60</sup>A | n.a.<sup>e</sup> | n.a. | n.a. | n.a. | n.a. | n.a. | 3 |
| F330<sup>6.52</sup>A | 4.4 | 8.36 ± 0.11 | 87 ± 4 | 3.2 | 8.50 ± 0.08 | 87 ± 3 | 3 |
| N333<sup>6.55</sup>A | n.a. | n.a. | n.a. | n.a. | n.a. | n.a. | 3 |
| R336<sup>6.58</sup>A | n.a. | n.a. | n.a. | n.a. | n.a. | n.a. | 3 |
| A343<sup>ECL3</sup>W | n.a. | n.a. | n.a. | n.a. | n.a. | n.a. | 3 |
| E344<sup>ECL3</sup>A | n.a. | n.a. | n.a. | n.a. | n.a. | n.a. | 3 |
| L347<sup>ECL3</sup>A | n.a. | n.a. | n.a. | n.a. | n.a. | n.a. | 3 |
| I352<sup>7.35</sup>A | n.a. | n.a. | n.a. | n.a. | n.a. | n.a. | 3 |

<sup>a</sup>Data shown are means ± S.E.M. from at least three independent experiments performed in triplicate. Source data are provided as a Source Data file.
<sup>b</sup>The span is defined as the window between the maximal <sup>125</sup>I-CCK-8 response (E<sub>max</sub>) and the vehicle (no ligand).
<sup>c</sup>Sample size; the number of independent experiments performed in triplicate.
<sup>d</sup>The IC<sub>50</sub> was not measured due to lack of interaction in the structure.
<sup>e</sup>Cannot be calculated due to poor binding of the corresponding mutant.

**Extended Data Table 5.** Effects of NN9056 and CCK-8 on IP1 accumulation in wild-type (WT) and mutant CCK_A_Rs.

| Mutant | NN9056 |  |  | CCK-8 |  |  | Expression<sup>d</sup><br>(% of WT) |
| --- | --- | --- | --- | --- | --- | --- | --- |
|  | EC<sub>50</sub><br>(nM)<sup>a</sup> | pEC<sub>50</sub> ±<br>S.E.M.<sup>b</sup> | Span<br>(% WT)<sup>c</sup> | EC<sub>50</sub><br>(nM)<sup>a</sup> | pEC<sub>50</sub> ±<br>S.E.M.<sup>b</sup> | Span<br>(% WT)<sup>c</sup> |  |
| Wild-type | 13 | 7.90 ± 0.05 | 100 | 4.1 | 8.38 ± 0.05 | 100 | 100 |
| N98<sup>2.61</sup>A | 28 | 7.56 ± 0.13 | 66 ± 4 | 56 | 7.25 ± 0.11 | 102 ± 5 | 222 ± 8 |
| K105<sup>ECL1</sup>A | 8.4 | 8.08 ± 0.09 | 112 ± 4 | 2.6 | 8.58 ± 0.12 | 92 ± 4 | 162 ± 43 |
| T117<sup>3.28</sup>A | 24 | 7.62 ± 0.11 | 77 ± 4 | 5.7 | 8.25 ± 0.12 | 63 ± 3 | 97 ± 6 |
| M121<sup>3.32</sup>A | 28 | 7.55 ± 0.18 | 35 ± 3 | 5.5 | 8.26 ± 0.15 | 65 ± 4 | 149 ± 8 |
| M121<sup>3.32</sup>V | 21 | 7.69 ± 0.60 | 15 ± 4 | 36 | 7.64 ± 0.36 | 40 ± 6 | 119 ± 13 |
| M121<sup>3.32</sup>L | 9 | 8.05 ± 0.46 | 21 ± 4 | 120 | 6.92 ± 0.87 | 20 ± 8 | 120 ± 7 |
| M121<sup>3.32</sup>I | 12 | 7.91 ± 0.18 | 54 ± 4 | 73 | 7.14 ± 0.24 | 63 ± 7 | 107 ± 21 |
| Y176<sup>4.60</sup>A | 5.3 | 8.28 ± 0.13 | 67 ± 3 | 39 | 7.41 ± 0.15 | 56 ± 4 | 41 ± 3 |
| R197<sup>ECL2</sup>A | 35 | 7.46 ± 0.10 | 115 ± 5 | 482 | 6.32 ± 0.15 | 106 ± 8 | 110 ± 16 |
| W326<sup>6.48</sup>A | 6.9 | 8.16 ± 0.16 | 47 ± 3 | 13 | 7.88 ± 0.23 | 38 ± 4 | 26 ± 4 |
| I329<sup>6.51</sup>A | 4.6 | 8.34 ± 0.13 | 63 ± 3 | 57 | 7.24 ± 0.17 | 71 ± 5 | 86 ± 4 |
| F330<sup>6.52</sup>A | 5.4 | 8.27 ± 0.19 | 39 ± 3 | 6.8 | 8.17 ± 0.11 | 52 ± 2 | 43 ± 4 |
| N333<sup>6.55</sup>A | 8.7 | 8.06 ± 0.07 | 91 ± 3 | 312 | 6.51 ± 0.09 | 79 ± 3 | 123 ± 20 |
| R336<sup>6.58</sup>A | 19 | 7.78 ± 0.19 | 61 ± 5 | 556 | 6.26 ± 0.19 | 61 ± 6 | 139 ± 7 |
| L347<sup>7.30</sup>A | 26 | 7.58 ± 0.14 | 59 ± 3 | 393 | 6.41 ± 0.12 | 64 ± 4 | 49 ± 8 |
| S348<sup>ECL3</sup>A | 8.2 | 8.09 ± 0.15 | 49 ± 3 | 74 | 7.13 ± 0.13 | 51 ± 3 | 112 ± 7 |
| I352<sup>7.35</sup>A | 10 | 7.99 ± 0.17 | 59 ± 4 | 77 | 7.12 ± 0.17 | 59 ± 5 | 134 ± 18 |

<sup>a</sup>EC<sub>50</sub> values were determined after 1 h stimulation by increasing concentrations of NN9056 or CCK-8.
<sup>b</sup>All values are means ± S.E.M. from at least three independent experiments performed in triplicate.
<sup>c</sup>The span is defined as the window between the maximal NN9056/CCK-8 response (E<sub>max</sub>) and the vehicle (no ligand).
<sup>d</sup>Protein expression levels of CCK<sub>A</sub>R constructs at the cell surface were determined in parallel by flow cytometry with an anti-FLAG antibody and reported as percent compared to the WT CCK<sub>A</sub>R from at least three independent measurements performed in duplicate.

**Extended Data Table 6.** Binding of gastrin-17 to wild-type (WT) and mutant CCK_B_Rs in competition with ^125^I-CCK-8.

| Mutant | IC<sub>50</sub> (nM) | pIC<sub>50</sub> ± S.E.M.<sup>a</sup> | Span<sup>b</sup><br>(% of WT) | N<sup>c</sup> |
| --- | --- | --- | --- | --- |
| Wild-type | 54.6 | 7.26 ± 0.06 | 100 | 5 |
| T111<sup>2.61</sup>A | 62.3 | 7.20 ± 0.14 | 71 ± 4 | 4 |
| T111<sup>2.61</sup>N | 32.2 | 7.50 ± 0.11 | 85 ± 4 | 4 |
| S131<sup>3.29</sup>A | 47.1 | 7.33 ± 0.15 | 86 ± 6 | 4 |
| M134<sup>3.32</sup>A | 78.0 | 7.11 ± 0.14 | 112 ± 7 | 4 |
| Y189<sup>4.60</sup>A | n.a.<sup>f</sup> | n.a. | n.a. | 4 |
| H207<sup>ECL2</sup>A | n.a. | n.a. | n.a. | 4 |
| S219<sup>5.39</sup>H | 47.6 | 7.32 ± 0.25 | 52 ± 6 | 4 |
| Y350<sup>6.52</sup>A | 42.7 | 7.37 ± 0.20 | 75 ± 7 | 4 |
| R356<sup>6.58</sup>A | n.a. | n.a. | n.a. | 4 |
| H364<sup>7.27</sup>A | 40.0 | 7.40 ± 0.22 | 67 ± 7 | 4 |
| L367<sup>7.30</sup>A | n.a. | n.a. | n.a. | 4 |
| S368<sup>7.31</sup>A | 6.5 | 8.19 ± 0.19 | 56 ± 5 | 3 |
| I372<sup>7.35</sup>A | n.a. | n.a. | n.a. | 4 |
| H376<sup>7.39</sup>A | n.a. | n.a. | n.a. | 5 |
| Y380<sup>7.43</sup>A | n.a. | n.a. | n.a. | 4 |

<sup>a</sup>Data shown are means ± S.E.M. from at least three independent experiments performed in triplicate. Source data are provided as a Source Data file.
<sup>b</sup>The span is defined as the window between the maximal <sup>125</sup>I-CCK-8 response (E<sub>max</sub>) and the vehicle (no ligand).
<sup>c</sup>Sample size; the number of independent experiments performed in triplicate.
<sup>f</sup>Cannot be calculated due to poor binding of corresponding mutant.

